# A developmental atlas of somatosensory diversification and maturation in the dorsal root ganglia by single-cell mass cytometry

**DOI:** 10.1101/2022.06.01.494445

**Authors:** Austin B. Keeler, Amy L. Van Deusen, Irene Cheng, Corey M. Williams, Sarah M Goggin, Ashley K. Hirt, Shayla A. Vradenburgh, Kristen I. Fread, Emily A. Puleo, Lucy Jin, Christopher D. Deppmann, Eli R. Zunder

## Abstract

Precisely controlled development of the somatosensory system is essential for detecting pain, itch, temperature, mechanical touch, and body position. To investigate the protein-level changes that occur during somatosensory development, we performed single-cell mass cytometry on dorsal root ganglia from C57/BL6 mice, with litter replicates collected daily from E11.5 to P4. Measuring nearly 3 million cells, we quantified 30 molecularly distinct somatosensory glial and 41 distinct neuronal states across all time points. Analysis of differentiation trajectories revealed rare cells that coexpress two or more Trk receptors and overexpress stem cell markers, suggesting that these neurotrophic factor receptors play a role in cell fate specification. Comparison to previous RNA-based studies identified substantial differences between many protein/mRNA pairs, demonstrating the importance of protein-level measurements to identify functional cell states. Overall, this study demonstrates that mass cytometry is a high-throughput, scalable platform to rapidly phenotype somatosensory tissues.

## Introduction

Somatosensory neurons residing in the dorsal root ganglia (DRG) transmit diverse sensory stimuli to the central nervous system (CNS), including mechanical pressure, changes in limb position, temperature, pain, and itch. Previous studies have identified up to 13 subpopulations of mature sensory neurons in the peripheral nervous system (PNS) by the first month of development in mice, and up to 18 subpopulations in adulthood^1–5^. While somatosensory neurons are relatively well characterized at maturity, many fundamental questions with respect to their development remain unresolved. In particular, intermediate progenitor cell types of the DRG remain poorly characterized, and the molecular profiles that control cell type specification have not been defined. Identifying the molecular trajectories and cell fate decisions that control DRG development promises to improve our understanding of sensory disorders with developmental components such as congenital insensitivity to pain with anhidrosis (CIPA) and autism spectrum disorder (ASD)^6–8^.

Previous efforts to monitor the diversification and maturation of somatosensory neurons have primarily relied either on microscopy, which detects a small number of proteins simultaneously, or single-cell RNA sequencing (scRNA-seq), which detects a large number of transcripts simultaneously. scRNA-seq and the related technique, single-nuclei RNA sequencing (snRNA-seq), have been applied to characterize the molecular diversity of cell types in a wide range of adult and developing neural tissues^9–13^, including the dorsal root ganglion^2, 3, 14–23^, but no study to date has measured every day of development across embryonic and postnatal timepoints. Such temporal resolution is essential if we are to determine precise lineages of both abundant and rare cell types that are responsible for somatosensory perception. In this study, we leverage the relatively high throughput (1×10^6^ cells/hour) single-cell analysis technique, mass cytometry, to exhaustively profile the composition of the DRG at every day of development from embryonic day E11.5 to postnatal day P4.

Mass cytometry is a flow cytometry variant that uses rare earth metal isotope-labeled antibodies and other affinity reagents to quantify the abundance of proteins and other biomolecules at the single-cell level^24^. Commercially available reagents permit over 40 molecular markers to be measured and quantified simultaneously in each cell, including cell surface receptors and intracellular signaling molecules^25^, transcription factors^26^, cell cycle status and proliferation state^27^, and cell viability^28^. Mass cytometry has been used previously to characterize gliomal cells and microglia in neural tissues^29–31^, but until now it has not been applied to neurons or other glial cell types in the CNS or PNS.

To investigate DRG development with mass cytometry, we developed a 41-antibody panel including key transcription factors, neurotrophic factor receptors, and other protein markers known to play a critical role in the specification and maturation of DRG cell types. We applied this panel to measure single-cell DRG samples from E11.5, shortly after the DRG have coalesced from migratory neural crest cells, to P4, when somatosensory neurons have innervated their peripheral targets and have begun to mature into distinct functional types^1^. With this approach, we identified and quantified the abundance of 30 molecularly distinct somatosensory glia and 41 somatosensory neuron subtypes across embryonic and postnatal development in the DRG.

The 41 somatosensory neuron subtypes we identify here show complementary overlap with postnatal DRG neurons previously identified by scRNA-seq^2, 5, 19^. However, a time course comparison reveals that mRNA transcript abundance does not accurately predict protein abundance, which is the best representation of a cell’s functional state.

Collectively, the findings presented in this study demonstrate for the first time that mass cytometry is a high-throughput, scalable platform for single-cell analysis of neural tissues such as the DRG.

## Results

### Characterization of DRG cell types by their protein expression signatures

To identify and characterize cell types in the DRG by their protein expression signatures, we first adapted mass cytometry methods for neural tissues. This involved optimizing tissue dissection and cell dissociation techniques (**Methods**) and developing a 41-antibody staining panel for specific neuronal and glial subtypes (**Supplementary Table 1**), including general markers of neuronal development (TuJ1, NeuN, MAP2, NeuroD1, CD24, Cux1, PGP9.5), glial development (Vimentin, Nestin, Sox10, BFABP, GFAP, cMet, CD9, Oligodendrocyte marker O4 (OligO4)), endothelia (CD31), vascular smooth muscle cells (VSMCs) (PDGFRɑ), and leukocytes (CD45). We also included markers known to play a critical role in the specification and maturation of DRG neuron types, including key transcription factors (Islet1, Runx3) and neurotrophic factor receptors (TrkA, TrkB, TrkC, Ret, p75NTR)^1^. Each antibody was conjugated to a unique rare earth metal isotope with the MAXPAR diethylenetriaminepentaacetic acid (DTPA) polymer^32^, and then titrated to identify its optimal staining concentration using known-positive and known-negative control cells (**Extended Data Fig. 1a,b**).

Mouse DRGs were collected at daily time points from E11.5 to P4 (**Fig. 1a**). For embryonic time points, DRGs from a single litter were pooled for cell dissociation and mass cytometry analysis. For postnatal timepoints, the pups were first separated by sex, and then DRGs were combined to generate a pooled male and female sample from each litter. At least two biological replicates (i.e. pooled litters) were used for each developmental time point. After dissection, the pooled DRGs were dissociated into a single-cell suspension^33^ and then briefly incubated with cisplatin as a non-cell permeant viability stain^28^, followed by paraformaldehyde (PFA) fixation and long-term storage at - 80°C (**Fig. 1a**).

**Fig. 1.**
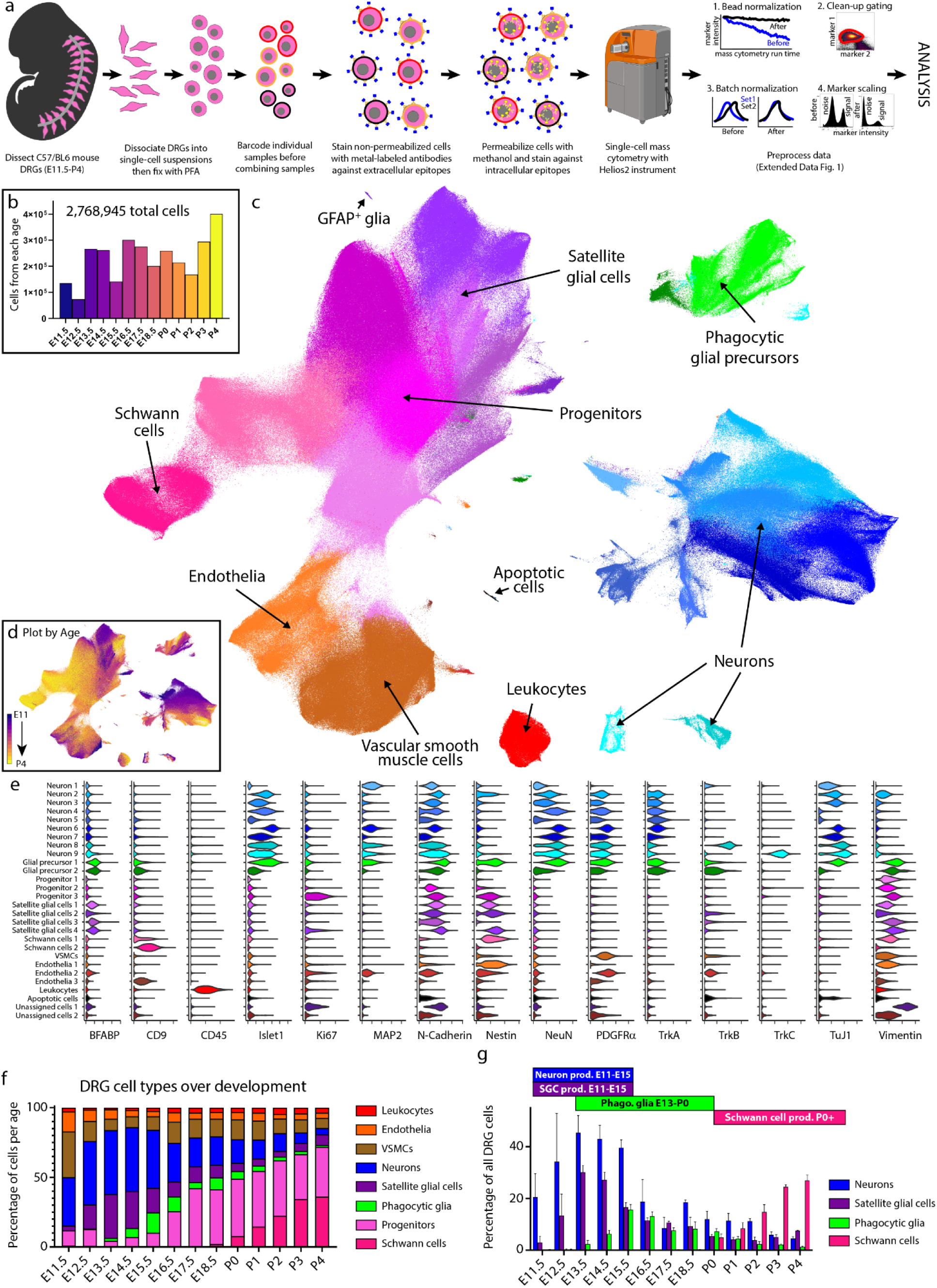
Characterization of DRG cell types from E11.5 to P4 by mass cytometry. **a)** Neural mass cytometry workflow. **b)** Total number of clean-up gated and preprocessed cells analyzed at each day of DRG development, consisting of 2,768,945 cells. These are from 2-4 separately analyzed litters (biological replicates) at each time point, for 43 samples in total. Bars are colored by time point. **c)** UMAP embedding of all DRG cells after clean-up gating, colored by primary Leiden clustering and labeled by presumptive cell type according to protein expression profiles. **d)** UMAP embedding from (c), colored by age. **e)** Violin plots of protein expression for clusters from c). **f)** The proportional abundance of major cell type classes across DRG development, created by combining the Leiden clusters from c) into 1) leukocytes: CD45^+^; 2) endothelia: CD31^+^, CD133^+^; 3) smooth muscle: p75NTR^+^, PDGFRɑ^+^, TrkB^+^, neuronal marker negative; 4) neurons: Islet1^+^, MAP2^+^, NeuN^+^, PGP9.5^+^, TuJ1^+^; 5) satellite glial cells: BFABP^+^, Sox10^+^, Vimentin^+^; 6) phagocytic glial precursors: BFABP^+^, Sox10^+^, Vimentin^+^, plus a mixture of neuronal markers such as Islet1^+^, MAP2^+^, NeuN^+^, PGP9.5^+^, TuJ1^+^; 7) neural progenitors: Ki67^+^, Nestin^+^, Sox10^+^, Vimentin^+^; and 8) Schwann cells: CD9^+^, cMet^+^, OligO4^+^. **g)** Changes in neuronal and glial abundance during periods of neuronal and SGC expansion, glial phagocytosis, and postnatal Schwann cell proliferation. Abbreviations: “phago.” stands for phagocytosis and “prod.” stands for production.

After all samples were collected and stored, they were thawed and barcode labeled^34^, followed by pooling into barcode sets, initially each containing a single replicate from E11.5 to P4 (**Supplementary Table 2**), for uniform staining and subsequent mass cytometry analysis (**Fig. 1a**). Preliminary inspection revealed that five samples had a low percentage of Islet1^+^ cells and two additional samples had low overall cell numbers, so additional litters were collected for these timepoints and run as a third barcode set (**Supplementary Table 3**). The resulting 4,974,302 events from three barcode sets were preprocessed by: 1) bead normalization (**Extended Data Fig. 2a**)^35^, 2) debarcoding^36^, 3) clean-up gating to remove dead cells, aggregates, and debris (**Extended Data Fig. 2b**; www.cytobank.org), 4) batch normalization^37^, 5) marker scaling (**Extended Data Fig. 2c**) and 6) removal of low complexity cells (**Extended Data Fig. 3a-f**). After these preprocessing and clean-up steps, we were left with 2,768,945 high-quality, viable, singlet DRG cells for analysis (**Fig. 1b; Supplementary Table 3**).

To identify broad classes of DRG cell types, we performed Leiden clustering^38^ on the full ∼2.8 million cell dataset and visualized the resulting clusters, or developmental age, on a 2D Uniform Manifold Approximation and Projection (UMAP) layout^39, 40^ (**Fig. 1c,d; Extended Data Figure 3g-j**). Cell type assignment for each cluster was determined by protein expression profiles matching neurons (23.09%), stem cells and glial progenitors (28.18%), satellite glial cells (12.21%), Schwann cells (11.87%), VSMCs (11.51%), endothelial cells (5.17%), and leukocytes (3.07%) (**Fig. 1e**). Because glial precursors can act as “non-professional” phagocytes^41^, we assigned the identity of putative phagocytosing glial precursors (4.9%) to cells that express glial precursor markers along with additional markers of their presumptive phagocytosed cargo (**Fig. 1e**). Our analysis to justify this classification is discussed further below.

To investigate how the abundances of all DRG cell types change across development, we grouped the cell populations identified by Leiden clustering (**Fig. 1c**) by cell class and calculated their abundance at each time point (**Fig. 1f and Extended Data Fig. 3k-z**). Generally, leukocytes, endothelia, and VSMCs showed consistent abundance across development. In agreement with previous studies, neurons expanded during early embryonic ages and then decreased in abundance, due to initial proliferation and migration waves, followed by a culling through competition for neurotrophic cues^42–47^. Consistent with previous studies, we observe that the relative abundance of neurons remains high until E16.5, at which point it diminishes due to the concurrent increase in glial progenitors (**Fig. 1f**)^43–47^. The proportion of neurons, satellite glial cells, putative phagocytic glial precursors, and Schwann cells across the time course correspond to the expected windows of neuronal proliferation and glial subtype expansion indicated in **Fig. 1g**.

### Comparison of DRG mass cytometry with immunohistochemistry

Neuronal cell types have not previously been characterized by mass cytometry, so we sought to validate our results by comparison with immunohistochemistry (IHC), assessing the total abundance of neurons observed with each technique, plus the relative abundance of TrkA, TrkB, TrkC, and Ret-expressing neurons. These neurotrophic factor receptors are essential for somatosensory growth and survival, but also delineate the broad neuronal cell types (**Fig. 2a**). While our previously described validation tests indicate good antibody specificity (**Extended Data Fig. 1**), we sought to determine whether specific cell types were selectively depleted during the mass cytometry processing steps and therefore underrepresented in our analysis. First, we performed IHC on DRGs from E11.5, E12.5, E13.5, E14.5, and P0 using Islet1 as a global marker for somatosensory neurons, assessed the percentage of Islet1^+^ cells out of total DAPI^+^ cells by IHC, and compared this to the percentage of Islet1^+^ cells observed by mass cytometry at the same ages, excluding non-DRG cells (blood, endothelia, VSMCs, Schwann cells) (**Fig. 2b**). In both our mass cytometry and IHC analysis, the percentage of Islet1^+^ cells was relatively consistent across E11.5 to E14.5, and then dropped approximately 50% by P0, which is due to both neuron death and expansion of non-neuronal cell types^43, 46, 47^.

**Fig. 2.**
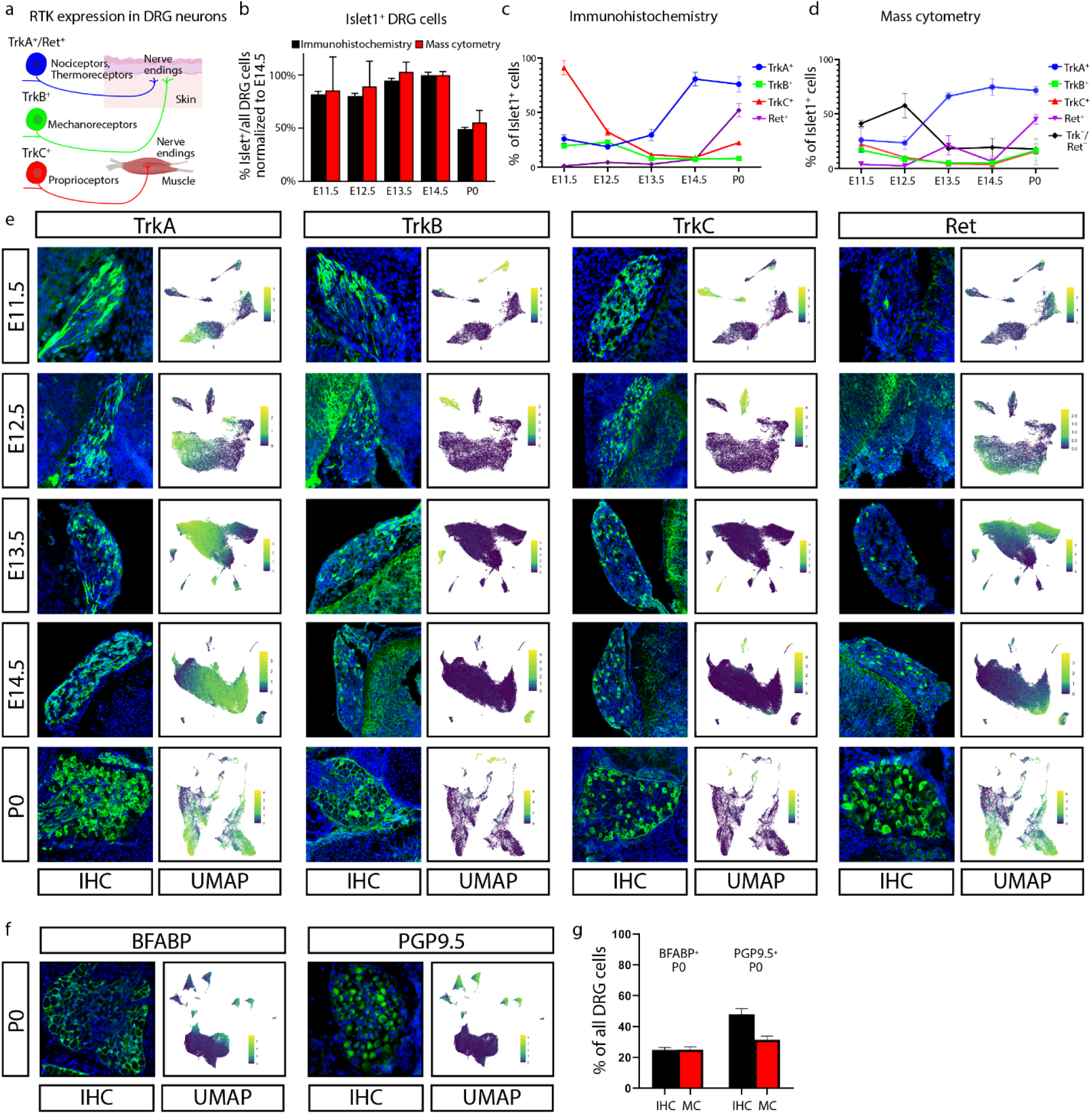
Comparison of DRG analysis by mass cytometry and IHC. **a)** Schematic illustration of somatosensory neuron subtypes divided by TrkA^+^, TrkB^+^, TrkC^+^, and Ret^+^ expression during development. Created with BioRender.com. **b)** Proportion of Islet1^+^ cells out of all DRG cells, by IHC and mass cytometry. All ages are normalized to E14.5, the peak of Islet1^+^ cell abundance by IHC. **c,d)** Proportion of TrkA^+^, TrkB^+^, TrkC^+^, and/or Ret^+^ neurons across matching timepoints between IHC (c) and mass cytometry (d), respectively. Mass cytometry also identifies the number of neurons that express Islet1 but none of TrkA^-^, TrB^-^, TrkC^-^, or Ret^-^. **e)** IHC of lower lumbar (L3-L6) DRGs stained for TrkA, TrkB, TrkC, or Ret; at ages E11.5, E12.5, E13.5, E14.5, and P0. IHC images are paired with mass cytometry UMAP layouts from neurons of the corresponding age, colored by protein expression for each RTK. **f)** IHC of P0 DRGs for BFABP and PGP9.5, and UMAP layouts of all P0 DRG cells colored by expression of these two markers. **g)** Relative abundance of BFABP^+^ and PGP9.5^+^ DRG cells at P0 by IHC or mass cytometry.

We next examined specific neuron subtypes by performing IHC on E11.5, E12.5, E13.5, E14.5, and P0 DRGs with antibodies against the receptor tyrosine kinases (RTKs) TrkA, TrkB, TrkC, and Ret, to distinguish the major classes of somatosensory neurons^1^. To compare cell abundance and staining intensity between mass cytometry and IHC, the microscopy images are shown side-by-side with UMAP plots from the same respective time points, subset for Islet1^+^ somatosensory neurons (**Fig. 2c-e and Extended Data Fig. 4a-c**). To quantify the abundance of somatosensory neuron subtypes observed by IHC, we counted the percentage of RTK^+^;DAPI^+^ double positive cells (**Fig. 2c**). To quantify the abundance of somatosensory neuron subtypes observed by mass cytometry the percentage of cells positive for each RTK was counted and then normalized to the percentage of Islet1^+^ cells from the same developmental age (**Fig. 2d**).

The patterns of cell abundance across this developmental window were similar for TrkA^+^, TrkB^+^, and Ret^+^ neurons, but the abundance of TrkC^+^ neurons differs substantially between IHC and mass cytometry at E11.5-E12.5 (**Fig. 2c,d**). Nearly all (∼90%) neurons express TrkC by IHC, but fewer (∼20%) do so by mass cytometry. It is possible that this could be due to preferential loss of TrkC^+^ neurons in these earliest embryonic samples during dissociation, cleavage of TrkC during early enzymatic cell dissociation, or detection of only cell surface TrkC in MC compared to the total TrkC detected in IHC.

We also compared IHC and mass cytometry with the glial marker BFABP and the neuronal marker PGP9.5 at P0 (**Fig. 2f,g**). At P0, the relative percentages of BFABP^+^ cells are comparable between techniques, although the proportion of PGP9.5-expressing cells is slightly lower by mass cytometry (**Fig. 2f,g**). This difference is likely due to the expansion of glial Schwann cell precursors in proximal nerve roots that can be spatially excluded by IHC but not by mass cytometry.

### Glial cell subtypes and their developmental trajectories

To further investigate how glia mature in the DRG, we selected all cell clusters that expressed glial markers Sox10, Vimentin, BFABP, CD9, cMet, OligO4, and GFAP; and then performed an additional round of Leiden clustering on this subset (**Fig. 3a, Extended Data Fig. 5a-d**), identifying four distinct glial cell types: 1) Schwann cells, 2) satellite glial cells (SGCs), 3) unspecified glial progenitors, and 4) putative phagocytic glial precursors. For a more detailed view of their molecular trajectories, we performed URD pseudotime analysis^48^, which uses timepoint-biased random walk iterations to identify the most likely cellular trajectory from a manually selected “root” cell type to manually selected “tip” cell types. For this analysis, we selected all E11.5 cells in the glial subset as the root, and each of the three mature cell types as tips (**Fig. 3b,c and Extended Data Fig. 5e-i**). From here, we performed an additional tertiary round of subclustering on the SGCs, phagocytic glial precursors, and Schwann cells (**Fig.** 3d,e,h,i,l,m).

**Fig. 3.**
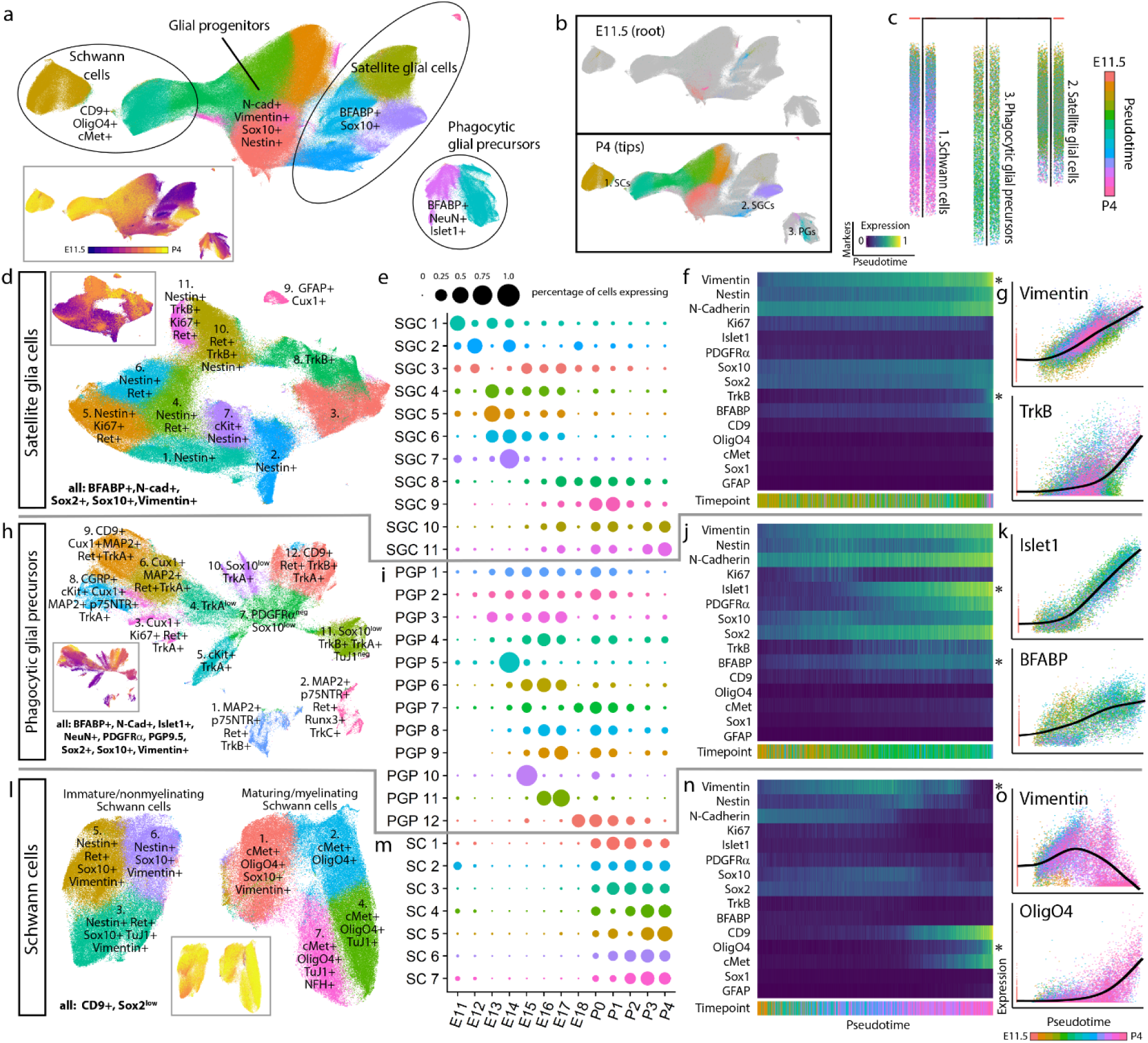
Glial subtypes show distinct developmental trajectories. **a)** Leiden clustering and UMAP embedding of E11.5-P4 DRG glia and glial precursors, performed on all cells from the clusters in Fig. 1c that expressed glial markers BFABP, CD9, cMet, GFAP, OligO4, Sox10, and/or Vimentin. General glial types were labeled along with key markers that delineated these cell types. Inset: colored by age. **b)** E11.5 or P4 cells only, colored by Leiden clustering, overlaid on the full UMAP embedding from (a), colored gray. **c)** URD pseudotime hierarchy for satellite glial cells, Schwann cells, and putative phagocytic glial precursors, produced with E11.5 root (pooled) and P4 tips (individual clusters) from (b), colored by pseudotime. **d-g)** Satellite glial cell clusters were extracted from the glial dataset for an additional secondary round of Leiden clustering and UMAP embedding, and SGC subtypes were labeled by their characteristic marker expression (d). Inset is colored by age. Relative abundances for each SGC subtype are plotted across sample collection ages (e), and changes in protein expression are plotted across the SGC pseudotime trajectory (f). Expression markers denoted by asterisks are also plotted by scatterplot (g) to illustrate the cells underlying the pseudotime heatmap in (f). Similar analysis was performed for putative phagocytic glial precursors **(h-k)** and Schwann cells **(l-o)**.

Glial precursors arrive in the DRG and start ensheathing somatosensory neuron cell bodies and phagocytosing dying neurons by E13.5^41, 49–52^. The non-myelinating, presumptive SGCs we observe here all express BFABP, N-cadherin, Sox2, Sox10, and Vimentin (**Fig. 3d-g**); and subcluster into eleven populations, four of which are retained at P4 (**Fig. 3d**). The early embryonic clusters express Nestin and a mix of Ki67, Ret, and c-Kit; while the postnatal SGCs upregulate TrkB (**Fig. 3d-g**). GFAP is a commonly used marker for SGCs, but only a small percentage of glial cells expressed GFAP, first appearing as ∼2% of all BFABP^+^ glia at E16.5, and increasing to ∼17% of BFABP^+^ glia by P4 (**Fig. 3d,e**).

Putative phagocytic glial precursors co-express non-myelinating glial markers (BFABP, Vimentin, and Sox10) but also contain neuronal markers (e.g. Islet1, NeuN, TuJ1) (**Fig. 3h-k**). Glial precursors have been shown to phagocytose dying neurons in the embryonic DRG before macrophages infiltrate and become the primary phagocytosing cell type^41^. We identified these putative phagocytic cells and performed the following analyses to rule out false positives associated with doublets and aberrant association with debris: 1) DNA intercalator gating, 2) “Barcode negative” filtering, 3) Abundance comparisons, 4) Expression intensity comparisons, and 5) Hemocytometer doublet analysis (discussed further in **Extended Data Fig. 2d,e** and **Extended Data Fig. 5k-n**). What are these glial precursors likely phagocytosing? Their subclustering is largely driven by the four RTKs that define somatosensory neuron types, TrkA, TrkB, TrkC, and Ret; but other neuronal proteins are also present, including nuclear transcription factors that indicate entire cell engulfment (**Fig. 3h, Extended Data Fig. 5m,n**). These putative phagocytosis events are observed across embryonic and postnatal development, but peak during the late embryonic stage (**Fig. 3i**).

Schwann cells (CD9^+^ and Sox2^low^) appear around birth and rapidly expand into two distinct classes: immature/non-myelinating (Nestin, Sox10, Vimentin) and maturing/myelinating (cMet, OligO4, and one subcluster still expressing Sox10 and Vimentin, suggesting a less mature state) (**Fig. 3l**). Each of these groups contain a subset of TuJ1^low^ cells, which is likely not expressed by these cells but rather comes from axons wrapped by these Schwann cells and not removed by dissociation (**Fig. 3l-o**). The neurofilament NFH is also present in a subset of the maturing/myelinating cells. These subtypes do not simply transition from the immature/non-myelinating (clusters 2, 4, and 5) to maturing/myelinating island (clusters 1, 3, 6, and 7), suggesting that the distinct islands are likely delineating between non-myelinating and myelinating pools of the same populations rather than a difference in developmental lineages or maturation state (**Fig. 3l,m**).

### Somatosensory neuron subtypes

To further investigate neuronal subtypes in the DRG, we extracted all clusters that expressed TuJ1, NeuN, Islet1, PGP9.5, and/or MAP2, and performed an additional round of clustering on this 637,744-cell subset. Specific subclusters were identified for removal as errant non-neurons, including putative phagocytic glia and spinal cord contamination (**Extended Data Fig. 6a**). After these cleanup steps, the remaining 533,488 cells were clustered again, dividing into three primary groups: TrkA^+^/Ret^+^, TrkB^+^, and TrkC^+^ (**Extended Data Fig. 6b-h**). These three groups were separated for a final round of subclustering to identify, as completely as possible, the somatosensory neuronal subtypes present across the DRG time course (**Fig. 4a-d**). From E11.5 to P4, we identified 41 neuronal cell types, including transient intermediates as well as all previously reported somatosensory neuron subtypes such as mechano-noxious heat peptidergic neurons (PEP), itch-mechano-heat non-peptidergic neurons (NP), cold-sensing neurons (remains a subpopulation within NP at P4, see **Extended Data Fig. 6i**), C-low-threshold mechanoreceptors (C-LTMR) (TH, see **Fig. 4f-n**), A-low-threshold mechanoreceptors (NF1, NF2, and NF3), and proprioceptors (NF4) (**Fig. 4d and Extended Data Fig. 6b-h**). These previously reported subtypes are labeled “E&E” in **Fig. 4d**, in reference to the review by Emery and Ernfors where they were described^5^.

**Fig. 4.**
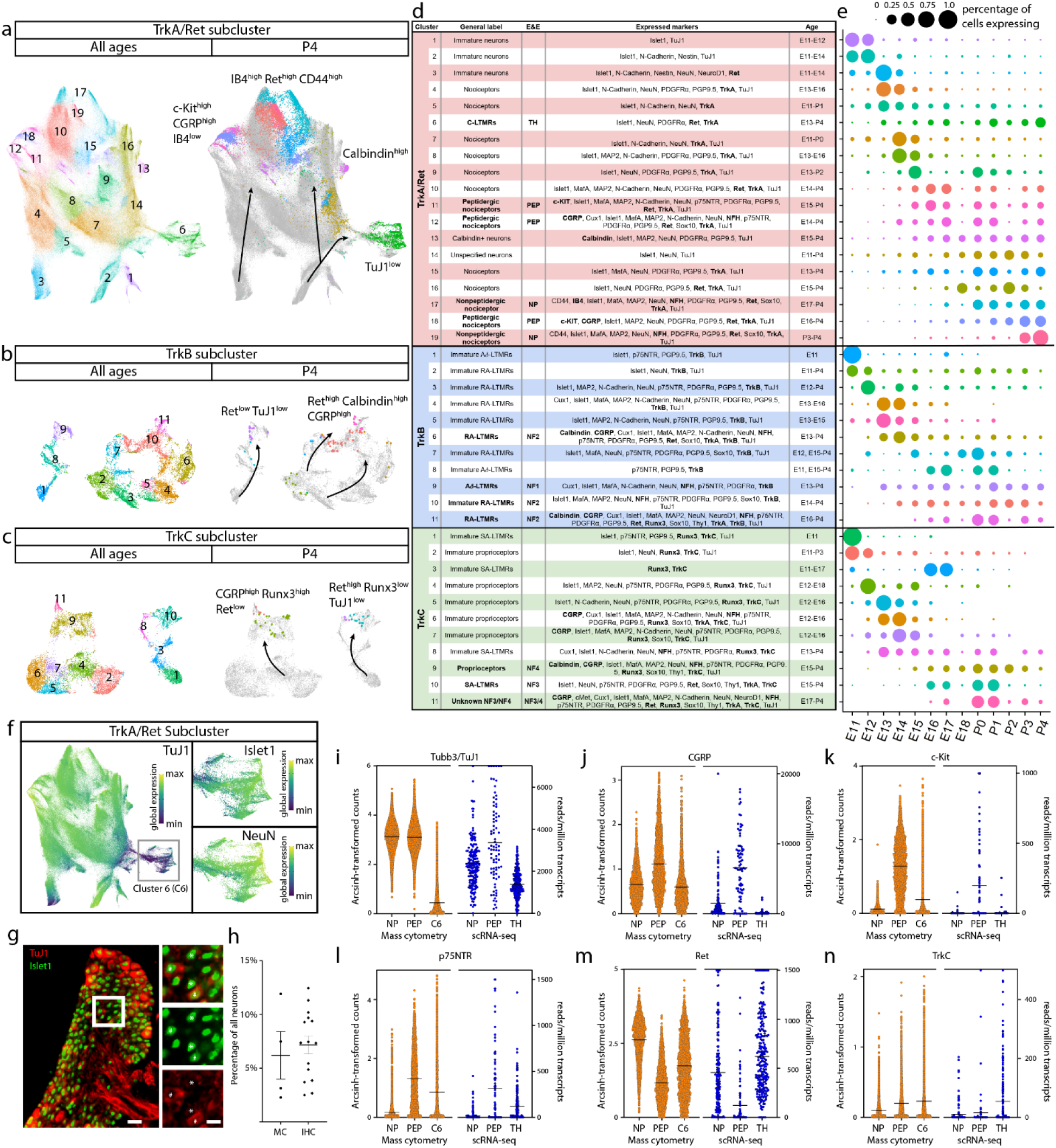
Distinct neuron subtypes emerge across development. **a-c)** Clusters from Fig. 1c expressing only neuronal markers (Islet1, MAP2, NeuN, PGP9.5, and TuJ1) were extracted for an additional secondary round of Leiden clustering and UMAP embedding (**Extended Data** Fig. 7), producing three distinct groups of somatosensory neurons characterized by TrkA/Ret, TrkB, or TrkC expression. Each of these three groups was extracted for an additional round of tertiary Leiden clustering and UMAP embedding for TrkA^+^/Ret^+^ **(a)**, TrkB^+^ **(b)**, and TrkC^+^ **(c)** neurons. Cells from all ages are shown in UMAP plots on the left, with clusters numbered by their age of first appearance. In UMAP plots on the right, all cells except P4 are colored gray to highlight the final time point populations. **d)** Table of all 41 neuron populations, listed in order of emergence by age. Cell types with protein expression that match designations from the recent Emery and Ernfors review are labeled^5^. Proteins expressed at a higher level relative to other neuronal populations are noted. Bolded proteins were used for subtype identification. The age range where each subtype was 1% or greater of all neurons in the TrkA/Ret, TrkB, or TrkC group is indicated. **e)** Relative abundance for neuronal subtypes, from E11.5-P4. **f)** TrkA/Ret UMAP embedding from (a) colored by TuJ1 expression. Cluster 6 with low TuJ1 expression is selected for additional visualization of Islet1 and NeuN expression in inset panels. **g)** IHC of P3 DRGs stained for Islet1 (Alexafluor488, green) and TuJ1 (Alexafluor568, red). Scale bar is 50 microns. Inset highlighting Islet1^+^ cells with corresponding high and low expression of TuJ1. Asterisk (*) denotes Islet1^+^;TuJ1^low^ neurons, pound sign (#) denotes Islet1^+^;TuJ1^+^ neurons. Inset scale bar is 20 microns. **h)** Percentage of TuJ1^low^ neurons observed by IHC (IHC, left) and mass cytometry (MC, right) at P3. For mass cytometry, two litters, sex separated were assessed (2 litters, 4 samples). For IHC, L4 DRGs were stained from 5 pups, 2-3 females and males per litter from 3 litters and counted for high or low expression of TuJ1 (3 litters, 15 DRGs in total). **i)** On the left, gene expression of Tubb3 (TuJ1) from the scRNA-seq dataset from Usoskin et al., 2015 (in blue) for nonpeptidergic nociceptors (NP), peptidergic nociceptors (PEP), and TH^+^ CLTMRs (TH) from P42-P56 mice^2^. On the right, protein expression of TuJ1 from mass cytometry for the same populations from P4 mice (in orange). TuJ1 expression was arcsinh transformed by 30. Means are denoted with a line. **j-n)** Comparison of mRNA transcript and protein expression from the same datasets for CGRP, c-Kit, p75NTR, Ret, and TrkC, respectively. Arcsinh transformation factors: 20, 20, 10, 20, and 15, respectively. Means are denoted with a line.

Separately clustering 493,544 TrkA^+^/Ret^+^ neurons, 21,652 TrkB^+^ neurons, and 18,292 TrkC^+^ neurons revealed 19, 11, and 11 distinct somatosensory subtypes, respectively, labeled here by their order of appearance during development (**Fig. 4e**). The TrkA^+^/Ret^+^ subtypes appear to emerge from three immature populations at E11.5 (clusters 1, 2, and 3) and diverge into four distinct subtypes: peptidergic nociceptors (c-Kit^+^, CGRP^+^, IB4^low^; clusters 11,12, and 18), nonpeptidergic nociceptors (IB4^+^, Ret^+^, CD44^+^; TrkA/Ret clusters 17 and 19), Calbindin^+^ neurons (TrkA/Ret cluster 13), and TH^+^ C-LTMRs (TuJ1^low^; TrkA/Ret cluster 6, see **Fig. 4f-n**) (**Fig. 4a,d**). The TrkB^+^ neurons separated into two distinct groups: Aδ low-threshold mechanoreceptors (Aδ-LTMR) (TrkB^+^, Ret^low^) and rapidly adapting low-threshold mechanoreceptors (RA-LTMR) (TrkB^+^, Ret^+^, Calbindin^+^, CGRP^+^). RA-LTMR clusters 2, 7, and 10 do not express the mature markers CGRP or Calbindin even at P4, indicating that these neurons do not mature until after P4 (**Fig. 4b,d**), although cluster 10 does express MAP2 and NFH. TrkC^+^ neurons separate into two distinct groups: slowly adapting low-threshold mechanoreceptor (SA-LTMR) (TrkC^+^, Ret^+^) and proprioceptors (TrkC^+^, Runx3^+^). Interestingly, a small population of mostly postnatal TrkC^+^ neurons (TrkC cluster 11) express both Ret and Runx3 (**Fig. 4c,d**), indicating that these cells may undergo a period of cell fate plasticity during innervation of their final tissue targets, between SA-LTMR and proprioceptors.

We were initially surprised to find neuronal clusters with low TuJ1 expression, as TuJ1 is a widely accepted marker of somatosensory neurons (**Fig. 4a-c**)^53–55^. TrkB^+^ and TrkC^+^ TuJ1^low^ clusters were identified by their differential expression of Ret and other key markers (Aδ-LTMR - TrkB^+^, Ret^low^, Calbindin^-^, CGRP^-^; SA-LTMR - TrkC^+^, Ret^+^, Runx3^-^). However, no markers were uniquely expressed in the TrkA^+^/Ret^+^ TuJ1^low^ cluster 6 (**Fig. 4f and Extended Data Fig. 6j-m**). To better understand these TuJ1^low^ neurons, we first sought to determine whether low TuJ1 expression was an artifact of incomplete mass cytometry staining or a bona fide somatosensory neuron subtype. We collected P3 L4/L5 DRGs and stained cryosections with TuJ1 and Islet1. These two neuronal markers were co-expressed in the majority of cells examined (**Fig. 4g**), but 7.16% (±0.8% SEM) of the Islet1^+^ cells had faint or no TuJ1 signal, compared to 6.2% (±1.9% SEM) by mass cytometry (**Fig. 4h**). In a publicly available scRNA-seq dataset on DRG neurons^2^, we identified a single TuJ1^low^ population, C-LTMRs (**Fig. 4i**), which exhibited a similar expression pattern (CGRP, c-Kit, p75NTR, Ret, and TrkC) to our TrkA^+^/Ret^+^ TuJ1^low^ cluster 6 (**Fig. 4j-n and Extended Data Fig. 6n**), and therefore, we labeled cluster 6 as TH^+^ C-LTMRs.

### Developmental trajectories of somatosensory neurons

The TrkA^+^/Ret^+^ and TrkB^+^/TrkC^+^ lineages are thought to be distinct as separate migration waves of specific progenitors, with a common progenitor in the neural crest^1^. Because this common progenitor is present prior to our earliest collection, we performed URD pseudotime analysis^48^ on each lineage separately, with unique roots for TrkA^+^/Ret^+^ (TrkA/Ret clusters 1, 2, and 3 at E11.5) vs. TrkB^+^/TrkC^+^ (TrkB clusters 1 and 2, TrkC clusters 1, 2, and 3, all at E11.5)(**Fig. 4a-d and Extended Data Fig. 7a-f**). Viewed together, these trajectories represent the maturation of 14 somatosensory cell types: mechanoreceptors (segments 1,4,7,9,14, and 17), proprioceptors (segments 10, 20, and 23), Th^+^ C-LTRMs (segment 6) peptidergic nociceptors (segments 11, 13, 15, 19, and 21) and non-peptidergic nociceptors (segments 12, 16, 18, and 22) (**Fig. 5a and Extended Data Fig. 7e-f**). As expected, these somatosensory subtype trajectories are distinguished by Trk paralog expression, as well as markers such as Ret, IB4, CGRP, and c-Kit. (**Fig. 5b and Extended Data Fig. 7g-t**).

**Fig. 5.**
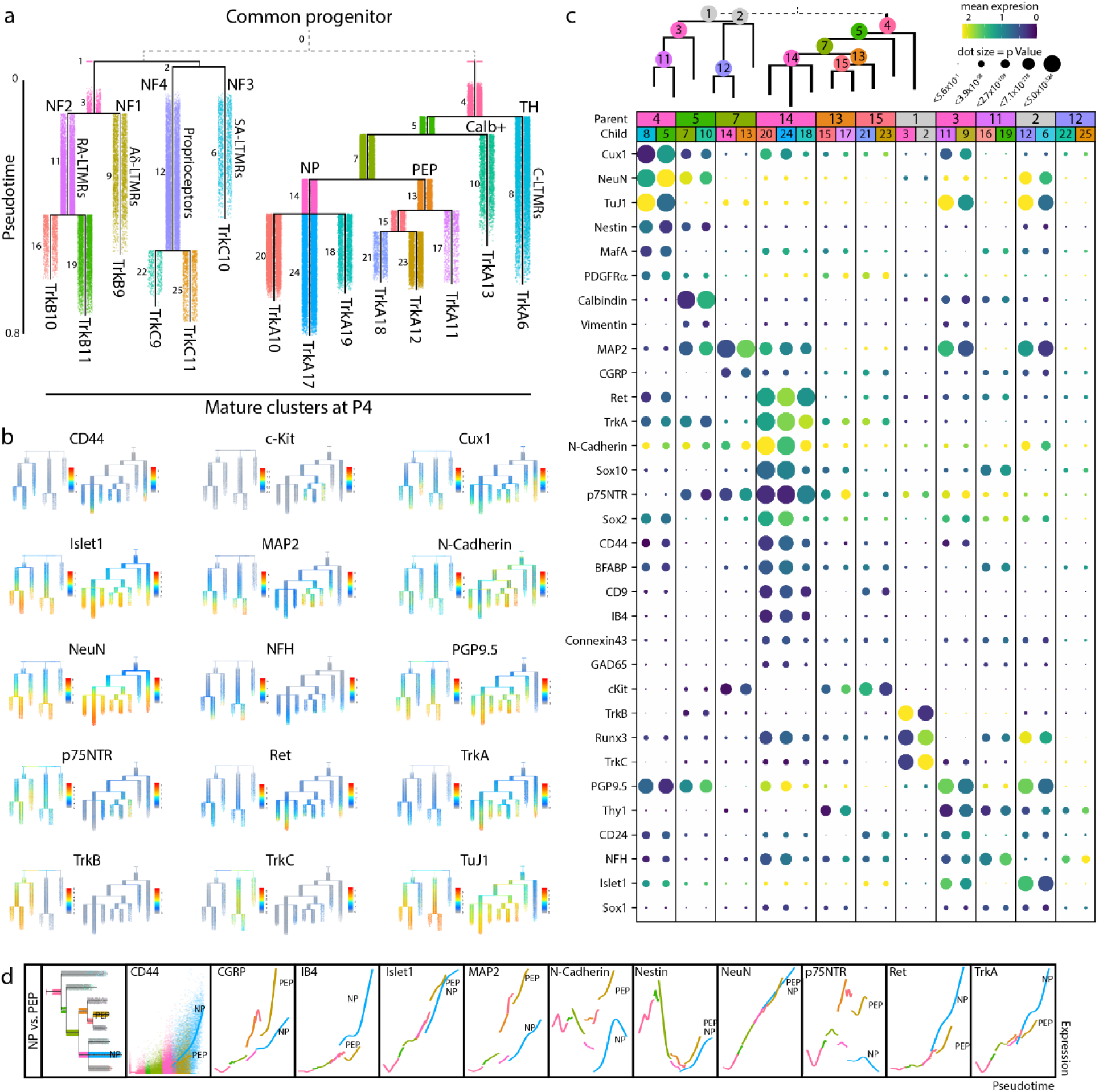
Neuronal differentiation tracked across development. **a)** URD pseudotime analysis of TrkB/TrkC neurons (39,944 cells) and TrkA/Ret neurons (64,997 cells downsampled from 493,544 total). These were run separately (grey dashed line), each with all E11.5 cells designated as root, and all mature P4 clusters designated as tips (**Extended Data Figure 9**). Cells are colored by their dendrogram segment, which are numbered by median pseudotime, from youngest to oldest. Key cell fate bifurcations are labeled at branch points. **b)** Proteins with the highest URD branch point divergence, ranked by p Value of the two-sample t-test between protein expression in the cell populations of each child segment. For branch points with more than two child segments, pairwise comparisons were made, and the most highly divergent p Value was used for rank ordering. Circle size indicates these p Values (log transformed) and circle color indicates marker expression in each child segment. **c)** Trajectory plots showing protein expression by pseudotime for NF2 (RA-LTMRs) and NF4 (proprioceptors). Each URD segment (colored in the grayed URD in the top left box in d) is graphed with all cells from that segment overlaid with a smoothed spline (e.g. CD24). **d)** Trajectory plots showing protein expression by pseudotime for nonpeptidergic and peptidergic nociceptors. Each URD segment (colored in the grayed URD in the top left box in e) is graphed with all cells from that segment overlaid with a smoothed spline (e.g. CD44). For clarity the dots representing the cells on both trajectories were removed from the subsequent markers.

To investigate cell fate decisions at URD branch points, we identified the most significant differences in protein expression between child segments at each branch point in the TrkBTrkC and TrkARet URDs (**Fig. 5c**). As expected, TrkB and TrkC are the key markers that divide TrkB^+^ RA-LTMR and Aδ-LTMR from the TrkC^+^ SA-LTMR and proprioceptors, along with Runx3 (**Fig. 5c**). Divergence between RA-LTMR and Aδ-LTMR trajectories (from parent segment 1 into child segments 7 and 9, see **Fig. 5a**) corresponds to changes in expression of Thy1, Cux1, Islet1, CD24, Calbindin, p75NTR, and NFH, among others (**Fig. 5c**). NeuN, Runx3, and N-Cadherin similarly show the largest shift in expression between SA-LTMRs and proprioceptors (**Fig. 5c**). In the TrkARet URD branchpoints, the split between peptidergic and non-peptidergic nociceptors corresponded to expression differences in MAP2, N-Cadherin, p75NTR, and c-Kit, suggesting that peptidergic nociceptors mature earlier than non-peptidergic nociceptors (**Fig. 5c**).

In fact, we observed differences in the timing and expression level of neuronal maturation markers across both pseudotime dendrograms. For example, MAP2 increases across all somatosensory populations except for the presumptive non-peptidergic nociceptors, indicating that maturation for this subtype is delayed until after P4 (**Fig. 5b,c**). The TNFR family member p75NTR, a pro-growth signal in somatosensory development, is expressed broadly in both TrkB^+^ and TrkC^+^ types, but only strongly in a single TrkA+ branch, a presumptive peptidergic (CGRP^+^,c-Kit^+^) cell type (**Fig. 5b,c**)^47^. CD44 signaling is involved in nociception and we observe early expression of CD44 in non-peptidergic nociceptors as well as the TuJ1^low^ TH cluster^56^ (**Fig. 5b,c**). Direct comparison of the peptidergic (PEP) and nonpeptidergic (NP) trajectories revealed an increase in CGRP, c-Kit, MAP2, N-Cadherin, and p75NTR for PEP, and elevated CD44, IB4, Ret and TrkA for NP (**Fig. 5d**).

### Double-Trk+ neurons exhibit elevated pro-growth and stemness markers

Since Trk receptors delineate somatosensory neuron subtypes and link a neuron’s survival to proper innervation of targets, we investigated whether neurons could transiently express two or more Trks simultaneously before committing to a specific cell fate. This analysis revealed a small pool of neurons that express two Trk receptors simultaneously (TrkA^+^;TrkB^+^, TrkA^+^;TrkC^+^, or TrkB^+^;TrkC^+^, respectively) (**Extended Data Fig. 8a-c**). These low frequency double-Trk^+^ neurons comprise less than 0.3% of the somatosensory neurons measured in this study, 1,480 out of the 533,488. Because they are so rare, most were not represented in our URD analysis, which was limited to approximately 60,000 cells by computational constraints on memory and runtime. To better investigate these rare cells, we mapped all double-Trk+ cells from our analysis onto the URD dendrograms by Euclidean distance, to estimate their trajectory and pseudotime position (**Fig. 6a and Extended Data Fig. 8d,e**). Interestingly, the incidence of these double-Trk^+^ neurons increases across pseudotime for many lineages (**Fig. 6a and Extended Data Fig. 8d,e**), and they exhibit characteristic shifts in the expression of pro-growth and stem cell markers when compared to single-Trk^+^ neurons from the same URD segment (**Fig. 6b**).

**Fig. 6.**
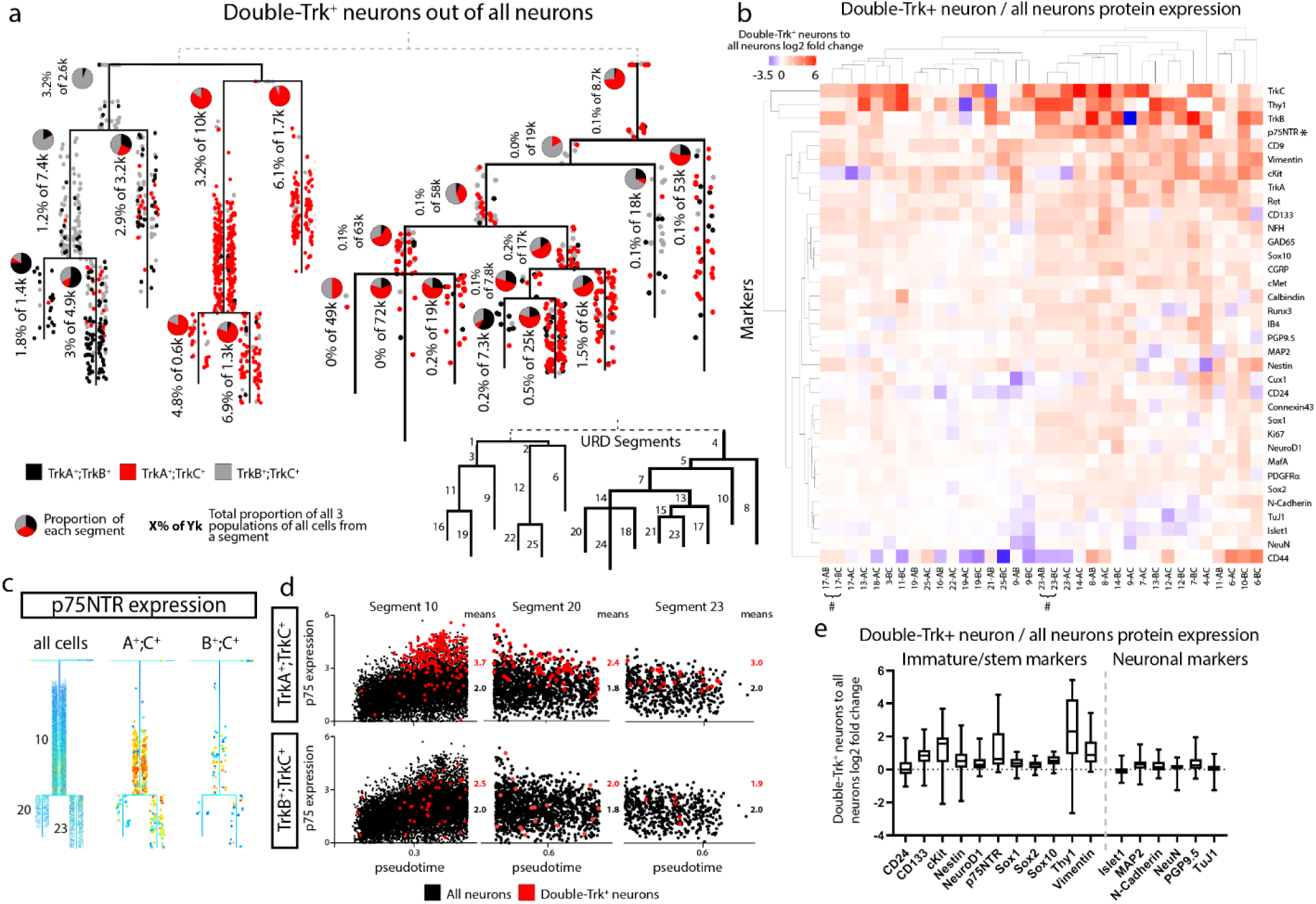
Elevated expression of stem cell and pro-growth markers in double-Trk^+^ neurons. **a)** 1,480 double-Trk^+^ neurons mapped onto the URD dendrogram from Fig. 5a by Nearest Neighbor Urd Trajectory Tool (NNUTT). All double-Trk^+^ cells on the TrkB/TrkC dendrogram come from the original URD analysis (no downsampling was used for URD construction), but the majority of double-Trk^+^ cells on the TrkA/Ret dendrogram were not in the original URD analysis (64,997 downsampled from 493,544 for URD construction). The relative proportions of TrkA^+^;TrkB^+^, TrkA^+^;TrkC^+^, and TrkB^+^;TrkC^+^ cells in each dendrogram segment are indicated by pie charts, and the total proportion for all double-Trk^+^ neurons relative to total neurons in each dendrogram segment is indicated as a percentage underneath each pie chart. **b)** Log2 fold-change of protein expression in the double-Trk^+^ neurons vs. total neurons in each URD dendrogram segment (Fig. 5a). Markers not expressed in neurons (e.g. GFAP, OligO4) were excluded from analysis. Segments were excluded from analysis with fewer than 10 cells and less than 0.1% proportion of double-Trk-expressor/all neurons, or less than 5 cells regardless of proportion. Pound sign (#) denotes paired segments where all double Trk neurons were, in fact, triple Trk neurons and the cells were the same cells producing identical expression profiles. **c)** URD dendrograms of the proprioceptor (segments 10, 20, and 23) population colored by expression of p75NTR for the full dataset and TrkA^+^;TrkC^+^ and TrkB^+^;TrkC^+^ populations of double Trk expressing neurons. **d)** Every cell in segments 10, 20, and 23, respectively, plotted by pseudotime value and p75NTR expression. All neurons colored in black with the double-Trk-expressors are overlaid in red. **e)** Marker expressions for immature/stem markers (left) and for neuronal markers (right) for all segments. Double-Trk-expressors were compared to all neurons of the same segment by log2 fold change.

The TNFR superfamily member p75NTR, a pro-growth receptor for somatosensory neurons^47, 57–60^, was expressed in TrkB^+^ and TrkC^+^ neurons from E11.5 to P4 and in peptidergic nociceptors, and not at all nonpeptidergic nociceptors (**Fig. 4d and Extended Data Fig. 6f,h**). In double-Trk^+^ neurons however, expression of the pro-growth p75NTR was elevated in many URD segments, such as TrkA^+^;TrkC^+^ neurons in the proprioceptor trajectory (**Fig. 6c,d**). Double-Trk^+^ neurons also showed increased expression of stemness markers but remained largely unchanged for other marker types (**Fig. 6e**). These rare cells could represent a state of increased plasticity before cell fate decisions are made. Alternatively, they may be neurons with delayed cell fate, or a residual pool of immature cells that is destined to act as neural stem cells in the adult DRG.

### Comparison of DRG mass cytometry with scRNA-seq

To compare our protein-based mass cytometry analysis with scRNA-seq, we selected the published study with the most closely overlapping time points: E11.5, E12.5, E15.5, P0, and P5 from Ginty and colleagues^19^. Our antibody panel has 36 proteins with directly comparable cognate mRNAs, but the other markers are not directly comparable, such as the lectin IB4 and cleaved Caspase 3. Because mRNA and protein levels are not strictly related in a linear or predictable manner (see analysis below), we first compared the normalized mean expressions and the percentage of cells expressing these 36 protein-mRNA cognate pairs. This comparison was performed at each matching time point, plus the close time points P4 (protein) and P5 (RNA) (**Fig. 7a**). In several cases, protein expression closely tracked RNA expression, such as Runx3, NeuroD1, N-Cadherin, TrkB, and TrkC (peak early); and NFH, CD9, CGRP, and Ret (peak late). In the majority of cases however, we observed large differences in the timing of relative expression levels between protein-mRNA cognate pairs. For example, the transcription factors Cux1, Islet1, MafA, NeuN, Sox1, Sox2, and Sox10 all exhibit peaks of relative protein expression that are delayed by at least 5 days relative to RNA expression, and we see similar patterns for BFABP, GAD65, PGP9.5, TrkA, cMet, CD133, c-Kit, MAP2, and GFAP. In rare cases, we even see relative protein expression levels that peak before RNA, such as for Thy1 and CD31.

**Fig. 7.**
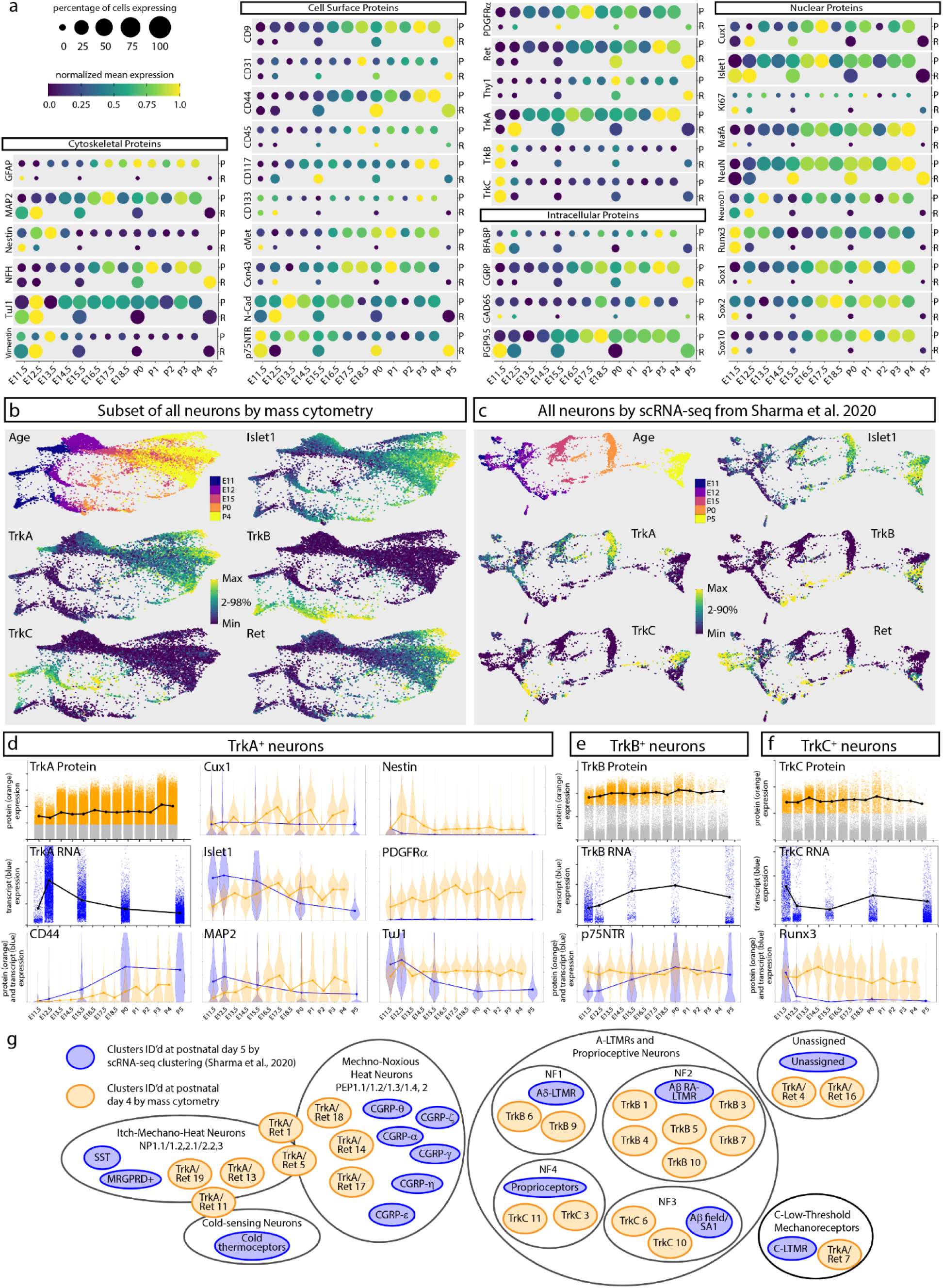
Comparison of DRG analysis by mass cytometry and scRNA-seq. **a)** Protein abundance measured by mass cytometry (E11.5 to P4) and mRNA transcript abundance measured by scRNA-seq (E11.5, E12.5, E15.5, P0, P5; Sharma et al., 2020^19^). Circle size indicates the percentage of cells with expression above marker-specific thresholds. For mass cytometry, the threshold was set at greater than the 99th percentile expression level of low complexity cells. For scRNA-seq, the threshold was set at zero; all non-zero values were included. Circle color indicates mean intensity of expression within the positive-expressing cells. Protein/mRNA pairs are grouped by protein subcellular localization. **b,c)** FLOW-MAP layouts from (a) mass cytometry (25,000 cells downsampled from all neuronal cells, Fig. 4a-c), and (b) scRNA-seq (full 32,169 cell dataset^19^), colored by age and marker expression level. **d-f)** Comparison of protein and mRNA expression levels in TrkA^+^, TrkB^+^, or TrkC^+^ neurons. Positive-expressing cells for each RTK were separated from non-expressers (gray) by thresholding for mass cytometry (orange) and scRNA-seq (blue) as described above. Violin plot overlays compare the normalized protein and mRNA abundance in each Trk-expressing population. **g)** Expression level overlap between the cell types identified by mass cytometry at P4 (orange) and scRNA-seq at P5^19^ (blue).

After comparing individual cognate pairs, we next sought to compare mass cytometry and scRNA-seq DRG measurements across development at the population level. Initial testing on the 32,169 neurons from the matched time points in Sharma et al.^19^ revealed that the gaps in developmental time between E15.5, P0, and P5 resulted in discontinuous Leiden clustering and UMAP embedding (data not shown), so the FLOW-MAP algorithm was applied to promote connections between the most similar cells from adjacent time points^61^. Along with the 32,169 scRNA-seq neurons, we analyzed approximately 25,000 mass cytometry neurons, evenly downsampled to approximately 5000 cells per time point (**Extended Data Fig. 9a**). By mass cytometry, three distinct trajectories correspond to TrkA/Ret (top), TrkB (bottom), and TrkC (middle) (**Fig 7b**); and by scRNA-seq, two distinct trajectories correspond to TrkA (top) and TrkB, TrkC, and Ret (bottom) (**Fig. 7c**). Islet1 expression increases similarly with age along each trajectory for both datasets. The observed differences in FLOW-MAP branching may result from RNA vs. protein differences (**Fig. 7a**), or the fact that our limited antibody panel was focused on somatosensory neuron development, based on functionally established molecular profiles^1^, as opposed to the transcriptome-level scRNA-seq measurements.

Because mRNA and protein levels are not strictly related in a linear or predictable manner, we cannot expect a 1-to-1 correspondence between mass cytometry and scRNA-seq cell clusters. As a simple approximation of this comparison however, we separated the TrkA^+^, TrkB^+^, or TrkC^+^ populations by thresholding (**Extended data Fig. 9b**), to compare RNA and protein expression in these cell types (**Fig. 7d-f**). As in the bulk comparison, we observe markers where RNA and protein levels track closely together, markers where the peak of RNA expression precedes the protein (Islet1 in all cell types, or CD44 in TrkA^+^ neurons), markers where the RNA levels swiftly diminish but the protein expression remains high (TuJ1 in all cell types, or Runx3 in TrkC^+^ neurons), and even markers where the peak of protein expression precedes the RNA levels (e.g. p75NTR in TrkB^+^ neurons).

While the cell types defined by clustering mass cytometry or scRNA-seq datasets are not expected to correspond perfectly, we were still interested to compare their similarities and differences. Because Ginty and colleagues found that early DRG timepoints were transcriptionally unspecialized^19^, we decided to focus on their P5 cell type clusters, and compare these with our P4 mass cytometry-based clusters. The two methods identified similar cell types, but interestingly, both methods found subpopulations not present in the other study (**Fig. 7g**). For example, while the scRNA-seq identified six CGRP subtypes, mass cytometry discerned at least three. Alternatively, while scRNA-seq identified a single cluster of Aβ RA-LTMRs, mass cytometry identified six molecularly distinct Aβ RA-LTMRs at P4. It is interesting to note that similar (yet complementary) results were obtained by mass cytometry with a panel of just 41 markers, compared to the transcriptome-level measurements by scRNA-seq.

## Discussion

Cell types in this study were identified by their protein expression profiles, using an antibody panel with canonical markers of somatosensory development^1^. The panel was limited to 41 antibodies by the number of available rare earth metal isotopes, and our antibody choices were limited by commercial availability. Although we did not have antibodies for every known marker of specific DRG subpopulations, our panel provided sufficient intersectional coverage to still identify these cell types. For example, while we were unable to validate an anti-TH antibody to use as a canonical marker of C-LTMR neurons, we were able to identify a combinatorial expression profile to identify C-LTMRs within our 41-antibody panel, even without anti-TH antibody. This combinatorial approach with non-canonical markers is made more powerful by quantifying expression levels within the linear range of mass cytometry, rather than simply categorizing them as positive or negative. Identifying the minimal set of markers that can be used to categorize every cell type in this way is an interesting algorithmic challenge, and an important consideration for mass cytometry in general. While there is always room for improvement, we consider the somatosensory-focused antibody panel presented in this study to be well-optimized for DRG phenotyping, and a useful template to build on for studies with a related molecular focus (**Fig. 8**).

**Fig. 8.**
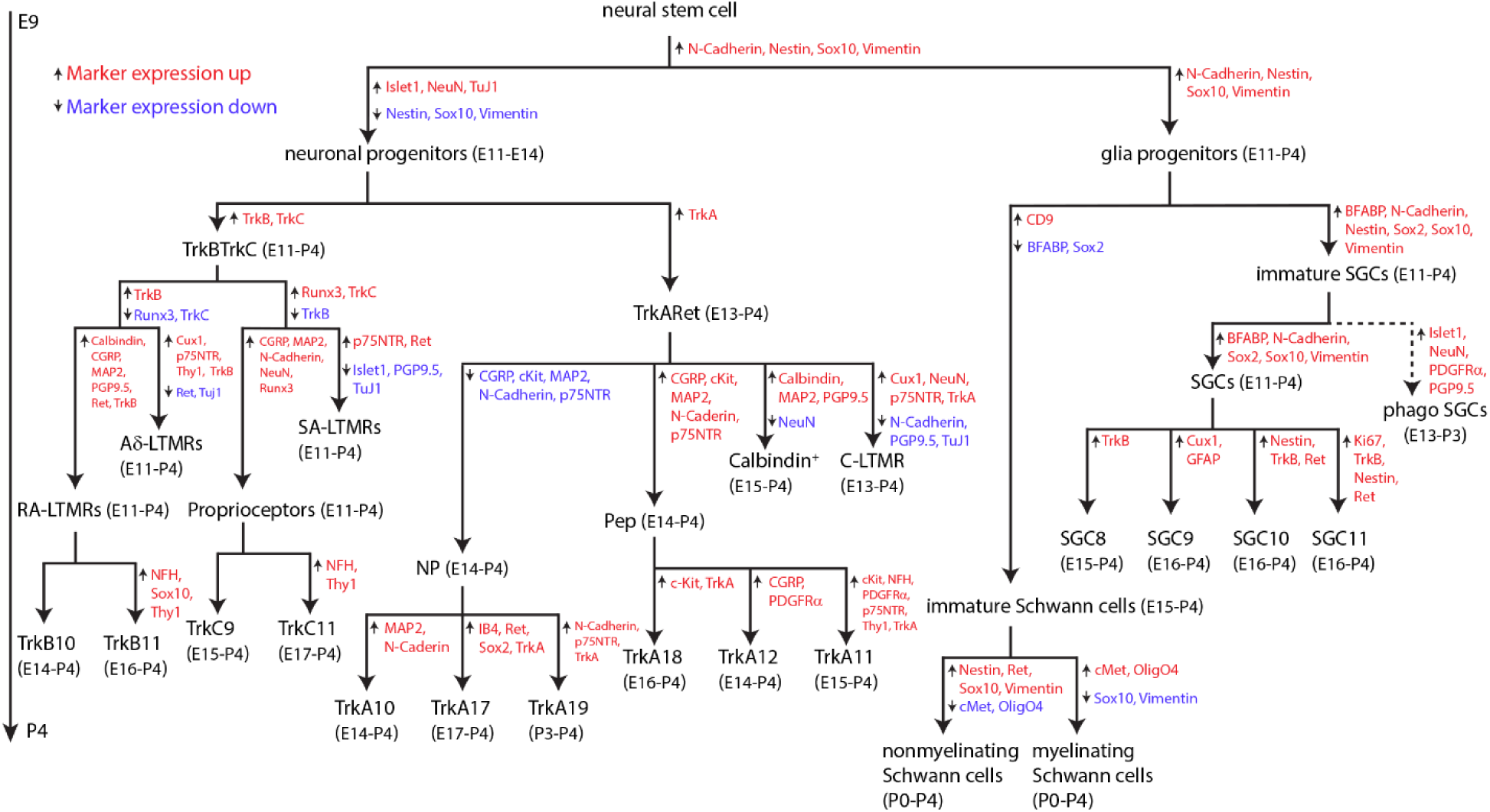
Somatosensory maturation and development in the DRG. Marker expression changes between segments are indicated by upward arrows with red text denoting marker expression increases and downward arrows with blue text denoting marker expression decreases. The age range where each population comprises at least 1% of that cell type (e.g. a neuronal cell type is at least 1% of all neurons, not all cells) is indicated beside each population label.

One related study of interest would be to examine cell signaling events in developing somatosensory cell types, to investigate which pathways contribute to cell fate decisions. This could be accomplished by combining a core set of identity marker antibodies from this study with a set of phospho-specific and other post-translational modification antibodies, to probe multiple cell signaling cascades simultaneously in every cell type. This approach has been applied to immune cells in suspension^62^, but cell dissociation presents a challenge because it can induce artifactual stress-related signaling. These effects can be mitigated by rapid dissociation and fixation from 2D culture^26^, or by fixing cells before dissociation from epithelial sheets^63, 64^ and organoids^65, 66^. DRG tissue is not compatible with these techniques due to the prolonged dissociation steps (Methods), but DRG explant cultures in vitro can be dissociated rapidly, and are therefore amenable to cell signaling analysis.

A prime candidate for cell signaling analysis are the rare double-Trk^+^ neurons we identified in this study, which are enriched for markers of stemness and proliferation. The double-Trk^+^ neurons may represent a transient cell state that can react to neurotrophic factors and switch cell fates if necessary. Previous studies have identified cell state-specific differentiation outcomes from the same neurotrophin stimulus^67^. The signaling mass cytometry approach would allow for the interrogation of these downstream pathways and cell states in every somatosensory cell type simultaneously, to determine how the internal response to neurotrophins such as NGF, NT-3, BDNF, and GDNF differs between cell types and contributes to cell fate and survival decisions. Another aspect of cell state, receptor internalization, could be added to this analysis by staining with one antibody before permeabilization, and the same antibody with a different metal label after permeabilization, to investigate the distinct roles of surface vs. internalized receptors^68^.

An unexpected result of this study was the observation of putative phagocytic events, consistent with previous reports^41^. We were not intending to investigate this phenomenon, so did not include glial subtype markers or markers of phagocytosis such as CD68^69^, Jedi-1, and MEGF10^41^ in our antibody staining panel. Based on our gating strategy (**Extended Data Figure 2b**), it is unlikely that the majority of putative phagocytic events result from cell doublets, larger aggregates, or incomplete dissociation of debris sticking to cells (**Extended Data Figure 5**). However, there were a few instances, particularly at later ages, where some of these events may have resulted from glial association with neurite debris from incomplete dissociation. Performing confocal immunofluorescence microscopy with markers of phagocytosis could help distinguish between these two possibilities. If the putative phagocytic events are corroborated, future studies could investigate whether glial subtypes have different patterns of phagocytic “food” over the course of development, and better characterize the molecular nature of these phagocytic events. This approach could provide additional insight into how the somatosensory nervous system is sculpted by axonal pruning and phagocytosis of whole cells and track the switch from phagocytosis by glial precursor cells to phagocytosis by macrophages later in development.

As illustrated in the discussion of putative phagocytic glia, an important caveat for single-cell analyses of neural tissue is that cell processes such as axons, dendrites, glial feet, and ensheathing membranes can be ripped away during tissue dissociation. For cell identification purposes, the loss of pre-synaptic markers is more problematic than post-synaptic markers, because the latter can also be detected in synapses on the cell body. Many axonal and dendritic proteins are synthesized in the cell body and can be detected here too, although at lower levels. In addition to marker loss, process shearing can also result in neurons appearing to express ensheathing membrane markers from glial cells, although neuronal identity is confirmed by the lack of glial nuclear proteins. In this case, the wrapping membranes can be viewed as an additional level of characterization for the neuron’s phenotype and maturity. In addition to process shearing, dissociation will introduce other cell state changes at the molecular level. As discussed above, cell signaling states are likely to be the most sensitive to perturbation, but proteolytic degradation and membrane protein internalization/trafficking can also change the molecular profile of dissociated cells from their original state in the tissue. All of these dissociation-related molecular changes are expected to be relatively consistent between mass cytometry experiments, facilitating direct comparison between samples, but corroboration by alternative methods such as IHC is important to confirm in-tissue molecular states (**Fig 2**).

Another caveat for single-cell analysis of neural tissue is the potential for biased loss of specific cell types during dissociation and processing. This could be due to more fragile cell types lysing during cell dissociation, or stickier cell types adhering to the inner walls of storage and processing tubes. Another potential source of biased cell loss is incomplete tissue dissociation, because the cells present in non-dissociated tissue are filtered out with a 40-micron strainer and lost to analysis. As above, IHC can be applied to validate the molecular profile of cell types and count their abundance in tissue slices (**Fig. 2**). Additionally, the same panel of metal-labeled antibodies developed for mass cytometry can also be applied to tissue sections for mass spectrometry-based imaging, by either imaging mass cytometry (IMC)^70^ or multiplexed ion beam imaging (MIBI)^71^. The ability to visualize DRG cells in their native tissue environment with the same antibody panel represents a parsimonious solution to the dissociation and processing-related caveats discussed above. Measuring cell morphology and localization with the same antibody panel presented in this study will yield additional insight, although cell segmentation is still a challenging problem. Mass cytometry still has the advantages of higher cell throughput, improved cell quantification/resolution, and comprehensive tissue coverage, as opposed to a single slice through the tissue.

After performing the first protein-level single-cell analysis of DRG by mass cytometry, we were interested to compare our results with published scRNA-seq studies. Expression profiles from scRNA-seq have previously been used to define DRG cell types, but mRNA expression cannot directly predict protein expression and functional cell states. For example, translational control, protein degradation, and incomplete trafficking or internalization of surface proteins may result in high levels of mRNA but no protein present. On the other hand, mRNA degradation may result in high levels of long-lived proteins, but no mRNA remaining present. The discrepancy between RNA and protein is thought to be largest during dynamic cell transitions^72, 73^, such as DRG development. We therefore sought to determine how well scRNA-seq could predict protein expression and functional cell states in the DRG.

While no scRNA-seq study covers every time point from E11.5 to P4, the best overlapping study to date is from Ginty and colleagues, covering E11.5, E12.5, E14.5, and E18.5, plus P5 which is closely adjacent to P4^19^. In our analysis, many protein-mRNA cognate pairs showed poor agreement across time points, highlighting the value of protein-level measurements by mass cytometry to identify functional cell states in the DRG. These differences could be regulated by translational control, protein and RNA degradation half-lives, protein trafficking for surface receptors, or any combination of these factors. The mechanisms that control these observed differences are likely to be different for each protein-mRNA cognate pair and could change considerably across developmental stages and between cell types. Uncovering and characterizing these mechanisms is beyond the scope of this study, but could be investigated further by parallel measurement of split samples with scRNA-seq and mass cytometry, and by performing CITE-SEQ to simultaneously detect protein and RNA at the single-cell level, although this technique is limited to cell surface proteins^74^.

Collectively, our high dimensional analyses of the DRG demonstrate replicable ground truths associated with sensory development. This technique provides inroads for future studies to ask fundamental developmental questions at an unparalleled pace and resolution.

## Methods

### Animals

All animal experiments were carried out in compliance with policies of the Association for Assessment of Laboratory Animal Care and approved by the University of Virginia Animal Care and Use Committee (Deppmann protocol no. 3795). Mice aged embryonic day 10.5 (E10.5) to postnatal day 4 (P4) were harvested from C57/BL6 females (Jackson Labs, 000664) bred in house. For timed pregnancies, animals were mated overnight and separated after 16 hours. Animals were housed on a 12-hour light/dark cycle with food and water ad libitum. For embryonic time points, pregnant females from single overnight (harem set up between 5-6pm and split between 7-8am the following morning) timed matings were used to ensure accurate embryo age.

### Dissection

Spinal cords were removed from mice aged embryonic day 11.5 (E11.5) to postnatal day 4 (P4) and placed in 35-mm Petri dishes containing Dulbecco’s phosphate-buffered saline (PBS; Thermo Fisher Scientific, 14190) on ice. DRG were plucked either off the isolated spinal cord (E11.5-E15.5) or from within the ossified vertebrae (E16.5-P4), depending on age. All DRG were collected upon dissection, including the sacral, lumbar, thoracic, and cervical ganglia. Total numbers of animals and cells analyzed are listed in **Supplementary Table 2**.

### Single-cell dissociation

After dissection, all the DRG from a whole litter (6+ pups) are transferred to a 15 ml conical filled with cold DMEM/F12. Excess media is then removed, followed by the addition of 5 ml of Enzyme Solution 1. For tissues E15.5 to P4, Enzyme Solution 1 is composed of 5 mg/ml bovine serum albumin (BSA; Sigma-Aldrich, A9418), 2 mg/ml Collagenase Type 2 (Worthington, LS004176), 0.2 mg/ml DNAse-I (Sigma-Aldrich, 11284932001), and 0.2 mg/ml hyaluronidase (Sigma-Aldrich, H3884) in DMEM/F12. For E11.5 to E14.5 tissues, Enzyme Solution 1 was prepared as above, and then diluted 1:10 in DMEM/F12. After 20 minutes of incubation in Enzyme Solution 1 at 37°C, this was then removed and replaced with 5 ml of Enzyme Solution 2 (Trypsin in DMEM/F12) for tissue from E15.5 to P4. Again, E11.5 to E14.5 was treated with Enzyme Solution 2 diluted 1:10 in DMEM/F12. The tissue was incubated for 15 minutes in Enzyme Solution 2 at 37°C before the solution was removed, leaving a residual volume of approximately 750 μl. Serial dissociation was performed with 4 fire polished pipettes with decreasing pore diameter, with approximately 10 triturations per pipette.

Our comparisons to IHC and scRNA-seq indicate that generally we were able to retain known cell types through our workflow and analysis. We also were not able to collect tissue older than P4 with enough efficiency and yield for mass cytometry analysis, due to the abundance of myelin and debris. Alternative dissociation methods or post-dissociation cleanup may make mass cytometry analysis possible for DRG past P4, including adult tissue.

### Cisplatin staining and fixation of cells

Final cell suspensions were mixed with 100 µl of 2✕ cisplatin solution (10 µM in PBS; Sigma Aldrich, P4394) and incubated for 30 sec before quenching with 1.3 ml of PBS containing 0.5% BSA. After centrifugation at 300 ✕ g for 3 minutes at 4°C, resulting cell pellets were washed once with PBS with 0.5% BSA before fixation in 1 ml of 1.6% paraformaldehyde solution (Electron Microscopy Services, CAS 30525-89-4) in PBS for 10 minutes at room temperature. After centrifugation at 600 ✕ g for 3 minutes at 4°C, cells were washed once with PBS before final resuspension in 1 ml of cell staining medium (CSM; 0.5% BSA, 0.02% NaN3 in PBS) and then the cell suspension was passed through a 75 µm sieve and 45 µm sieve (Thermo Fisher Scientific, 50871316 and 50871319) with a P1000 micropipette. Then the cells were stored at −80°C until all samples were collected.

### Cell counts and visual inspection by light microscopy

Fixed cells were thawed and then visually inspected in bright field mode at 4X, 10X, and 20X on an EVOS AMF4300 microscope (Thermo Fisher Scientific, Waltham, MA) to determine if excessive amounts of cell clumping or debris were present in samples. Samples that passed visual inspection were counted using a Bio-Rad TC20 Automated Cell Counter (Hercules, CA) to determine cell number (**Supplementary Table 2**).

### Metal conjugation of antibodies

Purified antibodies were conjugated to metals (listed in **Supplementary Table 1**) for mass cytometry analysis using MaxPAR antibody conjugation kits (Fluidigm) according to the manufacturer’s instructions. After labeling, antibodies were diluted at least 1:2 to a final concentration ranging from 0.05–0.4 mg/mL in Candor PBS Antibody Stabilization solution (Candor Bioscience GmbH) for long-term storage at 4°C.

### Validation of antibodies

After conjugation to a specific metal isotope, each antibody was titrated using a variety of cell samples and counterstains. Antibodies that generated signal in DNA intercalator-positive cells that also correlated with one or more positive counterstains, but were absent in cells with a negative counterstain, were considered to be specific and reliable. Ideal concentrations for the discernment of relative protein expression were determined by the concentration with the greatest separation between signal in positive controls compared to signal in negative controls while minimizing background. Optimal staining concentrations for each antibody are listed in **Supplementary Table 1**.

### Sample barcoding, staining, and intercalation

Cells were thawed on ice, pelleted by centrifugation at 600 ✕ g for 3 minutes at 4°C, and the supernatant was discarded. After washing once with CSM, cells were resuspended in 0.5 ml of cold saponin solution (0.02% in PBS) containing one of 20 specific combinations of 1 mM isothiocyanobenzyl EDTA-chelated palladium metals to barcode samples, as previously described^34, 36^. After incubation at room temperature for 15 minutes on a shaker at 800 rpm, tubes were centrifuged at 600 ✕ g for 3 minutes at 4°C, the supernatant was discarded, and the cell pellet was washed three times with CSM. At this point, individual samples were pooled into a total of three barcoded sets for antibody staining.

For staining of surface epitopes, cells were blocked in CSM containing 10% (v/v) normal donkey serum (Millipore, S30-100ML) for 30 minutes at room temperature. Next, primary antibodies indicated as “surface” in **Supplementary Table 1** were diluted in CSM and added to cells (100 ul staining volume per 1 x 106 cells), which were incubated on a shaker at 800 rpm for 30 minutes at room temperature. After incubation, tubes were centrifuged at 600 ✕ g for 3 minutes at 4°C, the supernatant was discarded, and the cell pellet was washed three times with CSM. For intracellular staining, cells were permeabilized by adding ice-cold 100% methanol to fill the tube and incubating on ice for 10 minutes with vortexing every 2 minutes. Next, tubes were centrifuged at 600 ✕ g for 3 minutes at 4°C, and the supernatant was discarded. After washing cells once with CSM, primary antibodies listed as “intracellular” in **Supplementary Table 1** were diluted in CSM and added to cells on a shaker at 800 rpm for 1 hour at room temperature. After incubation, tubes were centrifuged at 600 ✕ g for 3 minutes at 4°C, the supernatant was discarded, and cells were washed three times with CSM.

After primary antibody staining, cells were incubated in 1.6% PFA containing 0.1 uM Cell-ID Intercalator-Ir (201192, Fluidigm) for 15 minutes at room temperature on a shaker at 800 rpm, or overnight at 4°C without shaking. After intercalation, cells were washed once with CSM, once with water, once with 0.05% Tween-20 (in water), and again with water. Cells were pelleted by centrifugation at 600 ✕ g for 3 minutes at 4°C, then kept on ice until run on the mass cytometer.

### Mass cytometry

Immediately before analysis on a Helios CyTOF 2 System (Fluidigm Corporation), cells were resuspended in water (approximately 1 mL per 1 x 106 cells) containing 1:20 EQ Four Element Calibration Beads (Fluidigm) and passed through a 40-µm nylon mesh filter. Cells were analyzed in multiple runs at a rate of 500 cells per second or less.

### Normalization and debarcoding

To control for variations in signal sensitivity across individual runs on the mass cytometer, raw .FCS data files were first normalized using EQ Four Element Calibration Beads (https://github.com/nolanlab/bead-normalization)35. Next, normalized .FCS files from each run were concatenated for each sample set. Concatenated .FCS files were then debarcoded using software to deconvolute the 6-choose-3 Pd combinatorial barcode^34, 36^, permitting identification of individual samples (https://github.com/zunderlab/single-cell-debarcoder)36. A new parameter was added to the FCS files: bc_neg, which is the sum of the 3 Pd measurements expected to be zero based on the cell barcode deconvolution assignment. High values for this bc_neg parameter indicate that the cell event in question is likely to contain two or more cells, and this was used for an additional clean-up gating step below.

### Isolation of quality, single-cell events

To isolate single cells from fragments/debris and clumps of multiple cells, the normalized and debarcoded .FCS files described above were uploaded to Cytobank (https://community.cytobank.org) and clean-up gating was performed according to the strategy illustrated in **Extended Data Fig. 2b**. First, a secondary debarcoding clean-up process was performed by gating out events with a high bc_neg values, followed by gating out events with a low barcode separation distance and/or high Mahalanobis distance. Next, singlets and quality events were isolated by comparing the ion count length, center, and width parameters. Then, cells that were alive at the time of fixation were distinguished from debris and dissociation-damaged or destroyed cells with a gate using DNA-intercalator and cisplatin viability dye. Six unused channels (120Sn, 127I, 133Cs, 138Ba, 140Ce, and 208Pb) were identified in Cytobank to contain some background levels and events high in these channels were then gated out. Finally, the 4th sample set exhibited an elevated background (in nearly all channels, NeuN shown) in the last third of the runtime. These events were gated out with a time gate.

### Batch correction

While sets 1 and 2 were exposed to a single master mix of antibodies, they were run independently after antibody staining. Further, set 3 was run later. To account for possible batch-dependent effects, batch correction was performed after isolating single-cell events as described above. To correct signal intensities of individual markers for batch effects, debarcoded .FCS files were subjected to the batch adjustment process described in Schuyler et al. 2019 (https://github.com/CUHIMSR/CytofBatchAdjust)37. Specifically, signal intensities for the following antibodies were corrected at the 50th percentile because they yielded approximately Gaussian distributions and produced mean signals with variance greater than 1% for a universal sample (mixture of all ages examined) included in each barcoded set: TuJ1, cMet, Connexin43, Sox2, CD9, CD117, Nestin, Sox1, CD24, NFH, CD133, CGRP, NeuN, Sox10, Ki67, Vimentin, Thy1.2, TrkC, Runx3, N-Cadherin, GAD65, Calbindin, MAP2, TrkA, MafA, Islet1, PDGFRa, Ret, Cux1, BFABP, Cleaved Caspase 3, TrkB, PGP9.5, p75NTR, IB4. Mean signals for OligO4, CD31, and CD45 had 2%–3% variance and normal distributions with truncated lower tails. Batch correction of CD31 and CD45 markers at 65th and OligO4 85th percentiles was determined to be effective at reducing the variance of mean signal. As the variance of mean signal for GFAP or CD44 was less than 1%, these markers were not subjected to batch correction.

### Leiden clustering

Cells were partitioned with Leiden clustering (https://github.com/vtraag/leidenalg)38 to identify molecularly defined cell types, with the nearest neighbors parameter set to 100. In some cases, this resulted in a memory error on 240GB High Mem compute nodes, and nearest neighbors = 15 was used instead. To assess if clusters were homogenous and unimodal, we inspected violin plots of marker expression for each cluster. To improve the homogeneity of cell populations, cell types of interest were subjected to multiple rounds of Leiden clustering. For the first round of clustering (primary clustering) and all subsequent rounds of clustering (secondary, tertiary, etc.), all 41 expression markers were used for Leiden clustering analysis. Cell cluster identities were annotated by comparison to previously reported expression profiles of DRG cell types.

### UMAP dimensionality reduction

41-dimensional mass cytometry datasets (including all antibody markers), were embedded into 2 dimensions by uniform manifold approximation and projection (UMAP, https://github.com/lmcinnes/umap/archive/0.2.4.tar.gz) with the following parameters: nearest neighbors = 15, metric = euclidean, local connectivity = 1, components = 2, epochs = 1000^39^.

### Identification of developmental cell trajectories with URD

Code for the URD algorithm (https://github.com/farrellja/URD)48, originally designed to run on scRNA-seq datasets, was modified to interoperate with mass cytometry. Code including modified functions and a script to run the full analysis is included (see Code Availability). Datasets were downsampled proportionally (to ∼65,000 cells) to accommodate the computational demands of URD except for the TrkB;TrkC dataset in **Fig.7** which was smaller (39,944 cells) than the computational limits encountered. The URD parameters used were floodPseudotime n=500, minimum.cells.flooded = 2, and knn = 100. The sigma parameter was determined individually for each dataset with global auto-detection via Destiny in the URD pipeline.

### NNUTT (Nearest Neighbor Urd Trajectory Tool) for analysis of rare cell populations

NNUTT was created to position a selected subset of cells on a full URD dendrogram. The cells in this subset could either 1) be included in the approximately 60,000 cells used to generate the URD dendrogram, 2) come from the larger non-downsampled original dataset, or 3) come from a completely different dataset, measured separately. This tool simply maps each cell to its nearest neighbor in the 60,000 cell URD set by expression marker euclidean distance. Subsequent analyses can then be performed to identify which dendrogram segments are best represented by each mapped subset, and how protein expression in the mapped subsets compares to the original dendrogram cells for each segment.

### Tissue processing for IHC

Embryonic and postnatal mice were euthanized by decapitation. The lower lumbar spinal columns were dissected and fixed in 4% PFA overnight before cyroprotection in 30% sucrose in PBS for 2 days, all at 4°C. The tissue was subsequently embedded in OCT (VWR #25608-930) and then cryosectioned into 10-micron sections. The DRG from lower lumbar segments were analyzed at E11.5 and E12.5, immediately above the lower limb buds. The L4 DRG was analyzed at E13.5, E14.5, and P0, using the last rib as a landmark for T13.

### Immunostaining

Mounted sections were warmed to room temperature and washed with 1× PBS three times for 5 minutes each. Antigen retrieval was performed for all antibodies by microwave boiling slides/sections in sodium citrate buffer (10 mM sodium citrate, pH 6.0). Sections were cooled to room temperature, sodium citrate buffer was replaced, and sections were microwaved until boiling again. Sections were then rinsed three times with 1× PBS and incubated with blocking solution (0.2% Triton X-100, 3% normal donkey serum) for 1 hour at room temperature. Sections were incubated with primary antibodies diluted as detailed below in blocking solution overnight at 4°C. Sections were washed with 1× PBS three times for 5 minutes each, incubated with secondary antibodies for 1 hour at room temperature protected from light, and then washed with 1× PBS three times for 5 minutes each. Sections were mounted in Fluoromount-G with DAPI (SouthernBiotech). Primary antibodies used in this study: Goat Anti-TrkA (R&D, AF1056, 1:200 or 0.2ug/mL, RRID:AB_2283049), Goat Anti-TrkB (R&D, AF1494, 1:100 or 0.2ug/mL, RRID:AB_2155264), Goat Anti-TrkC (R&D, AF1404, 1:500, RRID:AB_2155412), Mouse Anti-Beta III Tubulin (TuJ1) (Covance, MMS-435P, 1:1000, RRID:AB_2313773), Mouse Anti-Islet1/2 (DSHB, 39.4D5, 1:100, RRID:AB_528173), Goat Anti-Ret (Neuromics, GT15002, 1:1000, RRID:AB_1622006), Rabbit Anti-Sox10 (Gift from S. Kucenas, 1:5,000), Rabbit Anti-BFABP (Gift from C. Birchmeier and T. Müller, 1:10,000, Kurtz et al., 1994). Secondary antibodies used in this study: Alexafluor 488 Donkey anti-mouse (ThermoFisher, A-21202, 1:500, RRID:AB_141607), Alexafluor 633 Donkey anti-goat (ThermoFisher, A-21082, 1:500, RRID:AB_141493), Alexafluor 568 Donkey anti-rabbit (ThermoFisher, A10042, 1:500, RRID:AB_2534017).

### Cell count quantification

DRGs were sectioned into fifths (five representative sets) and each section was collected and stained with the indicated marker. All tissue was imaged on the laser scanning confocal Zeiss 780 NLO at 20× resolution in z-stacks at 3 μm intervals for manual quantification in Fiji^75^. Cells expressing each marker were counted and compared to counts of all cells determined from counting DAPI stained nuclei. To determine the percentage of Islet1^+^ and RTK^+^ cells observed by mass cytometry at these time points, we set a threshold value for each protein determined by the 99th percentile of expression of the low complexity cells as a measure of background: Islet1 > 0.75, TrkA > 0.9, TrkB > 3, TrkC > 2, and Ret > 1.9.

### scRNA-seq data acquisition

Pre-processed and annotated scRNA-seq data from Sharma et al., 2020 was downloaded from the data browser provided with their publication (https://kleintools.hms.harvard.edu/tools/springViewer_1_6_dev.html?datasets/Sharma2 <u>019/all</u>)^19^. As our analysis was focussed on DRG development, we excluded the adult mouse timepoint (P40) from our analyses. For a per-feature comparison with our mass cytometry dataset, we selected the transcripts (from within those that had passed QC thresholds) that corresponded to our protein markers. The scRNA-seq dataset (External resource Table 1) from Usoskin et al., 2015 was downloaded from the additional supporting data for the manuscript (http://linnarssonlab.org/drg/)2.

### Comparison of mRNA vs. protein expression

To compare scRNA-seq and mass cytometry, neurons from both analyses were first selected by thresholding for positive neuronal marker expression. For the scRNA-seq data, expression above zero was considered as ‘expressing’. For mass cytometry data, which typically exhibits low background for each marker similar to any antibody based technique, thresholds for labelling cells as ‘expressing’ were determined by the 99th percentile marker expression of the low complexity cluster 6 (**see Extended Data Figure 3**). For both datasets, expression values following respective preprocessing were per-feature range-normalized to fall between 0 and 1.

### Visualization of high-dimensional data with FLOW-MAP

To incorporate time as a variable in inferring developmental trajectories of cell populations, graph structures incorporating the indicated time points from mass cytometry and scRNA-seq datasets were generated with FLOW-MAP^26, 61^. FLOW-MAP output (.graphml files) was visualized with Gephi software (http://www.gephi.org)^76^ and force-directed layout was performed with the ForceAtlas2 algorithm^77^. For visualization, node size was adjusted to indicate cell type abundance, and the “prevent overlap” option was selected in Gephi to ensure all graph indices remained visible in the layout.

### Population sorting and mRNA/protein comparison

For Trk^+^ population selection and comparison between scRNA-seq and mass cytometry datasets, both were thresholded on positive expression for TrkA, TrkB, or TrkC. For protein expression, we established TrkA, TrkB, or TrkC expression value by the 99th percentile of expression of the low complexity cells as a measure of background: TrkA > 0.9, TrkB > 3, and TrkC > 2, respectively. For RNA expression, we included any expression value above 0 for each Trk transcript. Cells were sorted *in silico* based on these per-marker expression thresholds to extract “positive” cells for the respective markers at each timepoint.

## Author Contributions

A.B.K., A.L.V., I.C., C.D.D., and E.R.Z. planned all experiments. A.B.K., I.C., and L.J., collected tissue and performed single-cell dissociations. A.L.V. validated all antibodies in the DRG panel. A.B.K. and E.R.Z. performed the mass cytometry measurements. A.B.K., A.L.V., C.M.W, S.M.G., A.K.H., K.I.F., E.A.P., and E.R.Z. wrote the scripts for the analysis pipeline. A.B.K. performed data analysis. S.M.G. conceived of and performed the comparisons with scRNA-seq. E.R.Z. and C.D.D. conceived and supervised all aspects of the project. A.B.K., C.D.D. and E.R.Z. prepared figures and A.B.K., I.C., C.D.D. and E.R.Z. wrote the manuscript with input from all authors.

## Acknowledgments

Research reported in this publication was supported by the National Institute of Neurological Disorders and Stroke of the National Institutes of Health under award number F32NS103770 to A.B. Keeler, and R01NS111220 to E.R. Zunder and C.D. Deppmann. The content is solely the responsibility of the authors and does not necessarily represent the official views of the National Institutes of Health. Further support was provided by the 3 Cavaliers Pilot research program (to C.D. Deppman and E.R. Zunder). We thank N. Sharma and D.D. Ginty for consultation about data formatting and analysis steps from their scRNA-seq dataset. We thank Ali Güler, Amrita Pathak, Yipkin Calhan, Sushanth Kumar, Ekaterina Stepanova, Sonia Chandra, and Micah Hunter-Chang for feedback and critical evaluation of the data and manuscript. We thank the University of Virginia Flow Cytometry Core, RRID: SCR_017829 for technical assistance. The authors acknowledge Research Computing at The University of Virginia for providing computational resources and technical support that have contributed to the results reported within this publication. URL: https://rc.virginia.edu.

## Data availability

Further information and requests for resources and reagents should be directed to and will be fulfilled by Eli Zunder. The unprocessed single-cell mass cytometry datasets are accessible at FlowRepository (https://flowrepository.org/) [ID: FR-FCM-Z4S9]. The debarcoded sample FCS files and clean-up gates used for pre-processing are available at Cytobank (https://community.cytobank.org/cytobank/experiments/102249).

## Code availability

The code used to perform analysis of mass cytometry data was adapted from standard R and Python packages, including UMAP, LeidenAlg, and URD. The code is available on GitHub at https://github.com/zunderlab/Keeler-et-al.-DRG-Development-Manuscript. More detailed information is available upon request.

## Contact for reagent and resource sharing

Further information and requests for resources and reagents should be directed to and will be fulfilled by Eli Zunder.

---

**Extended Data Fig. 1.**
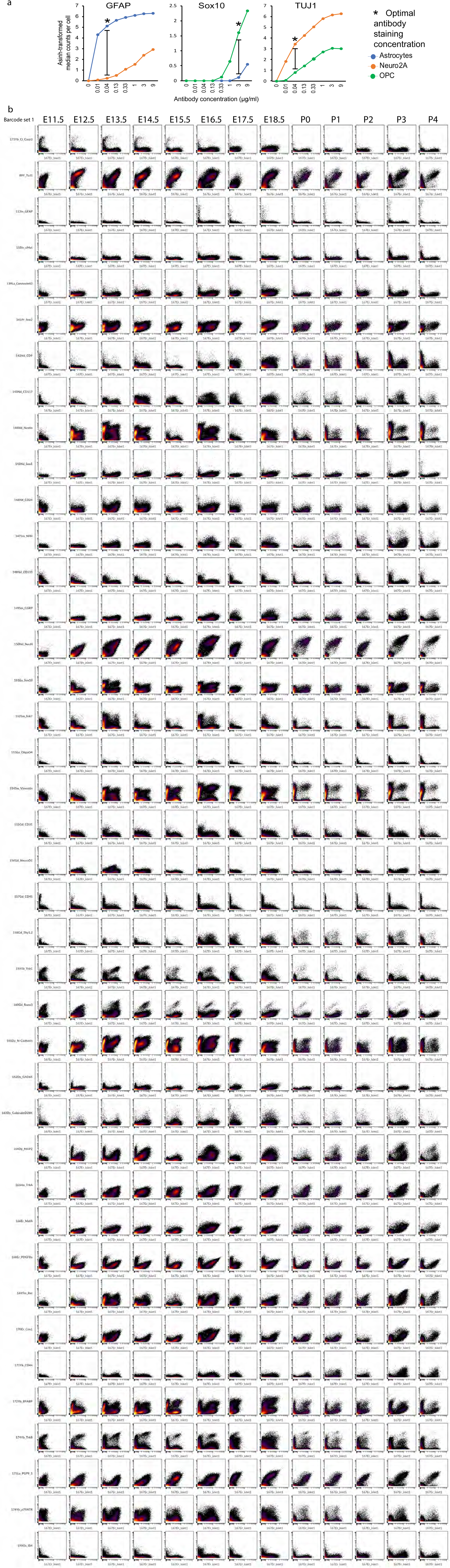
Validation of antibodies for mass cytometry. All antibodies validated and included in the DRG mass cytometry antibody panel. **a)** Each antibody was titrated across a range of concentrations (e.g. 9ug/ml to 0.01ug/ml). Known-positive and known-negative control cell samples were tailored for each antibody. Sometimes these were separate samples, and sometimes the known-positive and known-negative cells coexisted in a single sample, distinguishable by a separate antibody counterstain. Optimal staining concentrations for each antibody were determined by identifying the largest difference in signal intensity between known-positive and known-negative cells. **a)** Biaxials scatterplots for each antibody in the panel, demonstrating positive and negative staining across the DRG developmental time course. Full data available in Data Availability.

**Extended Data Fig. 2.**
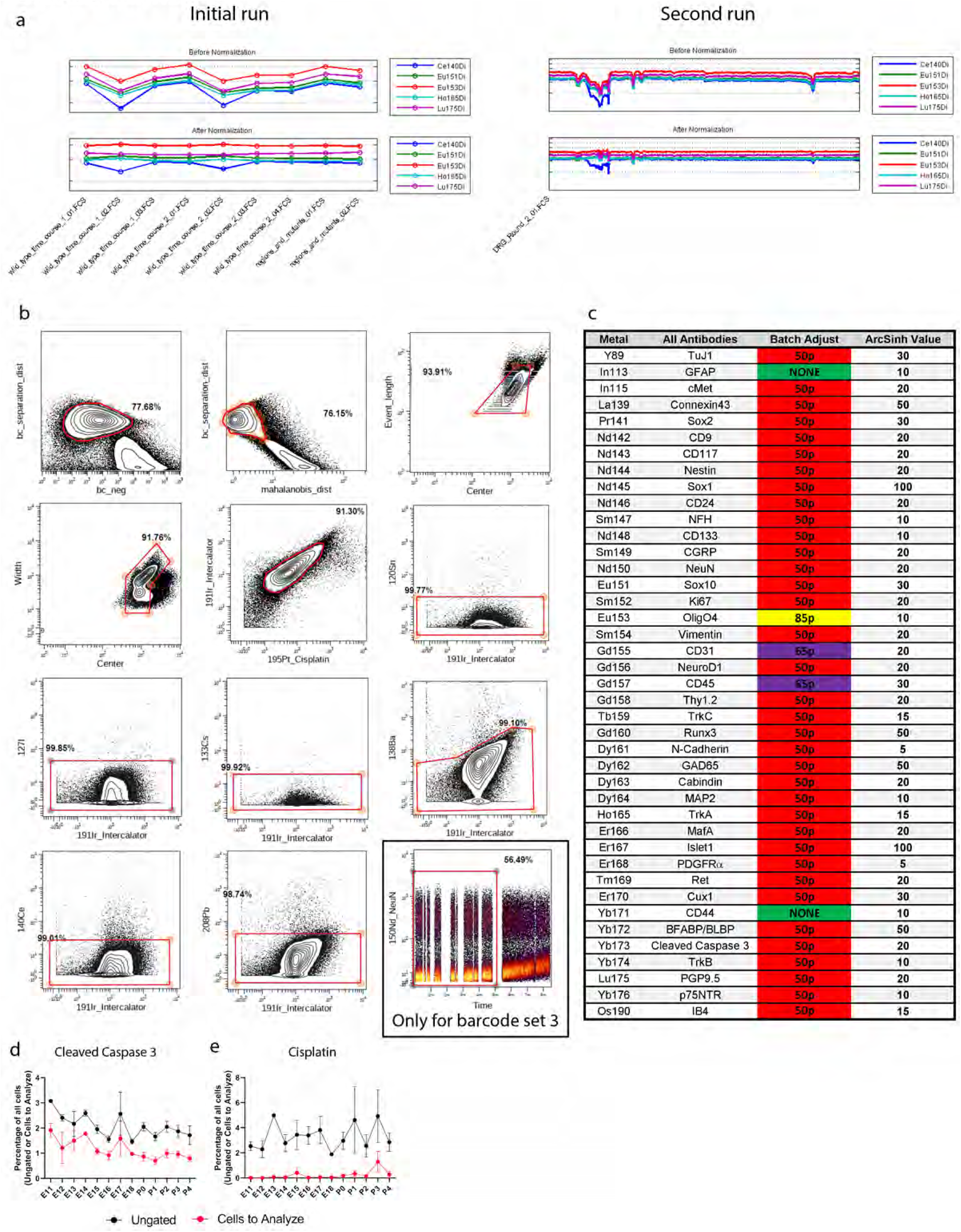
Pre-processing of DRG samples for mass cytometry. **a)** Calibration bead normalization of the raw mass cytometry data (stored as .fcs files) using the Matlab software described in Finck. et al., 2013^35^. **b)** Clean up gating done with Cytobank (cytobank.org) to remove low quality events from the dataset. Biaxial gates as follows: 1) barcode_separation x barcode_negative and 2) barcode_separation x malahoidis_distance removes events that cannot be confidently separated by barcode label; 3-4) event_length x center and width x center remove events that fall outside of normal Gaussian parameter distribution; 5) intercalator x cisplatin removes both non-cell events (e.g. cellular debris) and dying cells; 6-11) unused metals x intercalator removes high background events. A twelfth clean up gate was required for samples from barcode set 3 to remove a runtime-dependent increase in background in a subset of channels: time x NeuN. **c)** Batch correction was run to normalize signal strength between runs (Schuyler et al., 2019)^37^. Each barcode set included a “universal” sample consisting of excess samples from across the DRG time course. These excess cells were pooled together, and then aliquoted and stored at −80°C, to be included with each mass cytometry run as an unvarying control. After all samples were run, the universal samples between barcode sets were batch corrected to be as similar as possible on a per-marker basis, and then the batch adjustment process corrected the rest of the samples in that barcode set based on its corresponding universal sample. Arcsinh transformation values were manually adjusted to provide the greatest contrast between background and physiological values. **d)** Analysis of cleaved caspase 3 signal across the time course in the ungated sample (black line) and the cleanup-gated sample (red line). We see very little cleaved caspase 3 signal in the time course except for ∼E11.5-E13.5 in a small neuronal population. Here we see a small (∼2%) reactivity to cleaved caspase 3 in the ungated population that is mostly excluded in the cleanup gating. **e)** Analysis of cisplatin signal across the time course in the ungated sample (black line) and cleanup-gated sample (red line). Approximately 4% of the ungated events showed cisplatin staining, all excluded in the cleanup gating.

**Extended Data Fig. 3.**
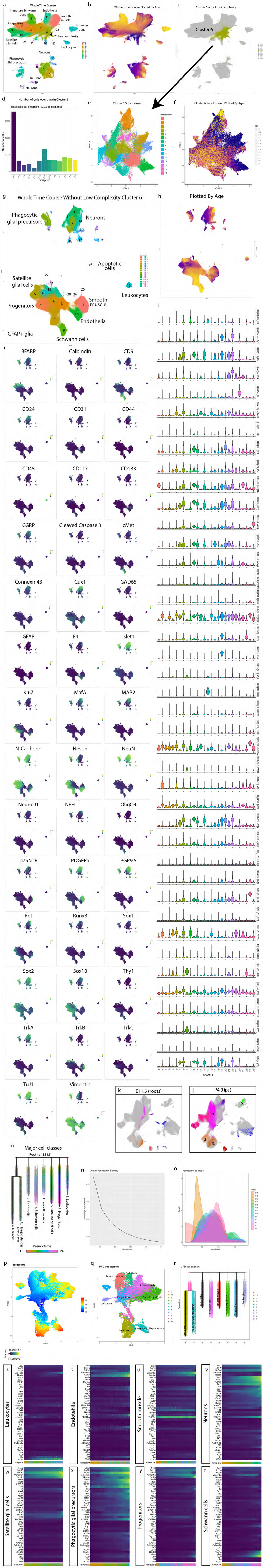
High dimensional analysis of the entire somatosensory time course data set. **a)** UMAP embedding of all Cells_to_Analyze (cleanup gating applied) from the whole DRG time course. **b)** UMAP plot from (a) colored by age. **c)** UMAP plot grayed out except for low complexity Cluster 6. **d)** Cells from all ages in Cluster 6, with predominant contribution from E11.5. **e)** UMAP embedding of Cluster 6 after extraction and secondary Leiden clustering. **f)** UMAP plot from (e), colored by age. **g)** Unmodified UMAP embedding from Fig. 1b. UMAP layout was rotated and white space removed for improved visualization. **h)** UMAP plot from (g) colored by age. **i)** UMAP plot from (g) colored by expression level for every marker in the DRG antibody panel. **j)** Violin plots of all markers for all clusters; Fig. 1e is truncated to show just the most salient markers for the general populations. **k,l)** Grayed out UMAP plots from Fig. 1c with only E11.5 (k) or P4 (l) cells superimposed on top, colored by general cell type as in Fig. 1f and Fig. 1g, respectively. A subsample of the cells from E11.5 were designated as the “root” for the subsequent URD analysis. Each cluster at P4 was subsampled and set as “tips’’ for URD. **m)** URD pseudotime dendrogram of the eight cell types to provide a molecular trajectory over maturation. **n)** During pseudotime calculation, several simulations are run, allowing pseudotime to be calculated from these iterations. For ideal pseudotime stability (e.g. decreased change in cell pseudotime with increasing runs) we assessed the number of runs required to approach an asymptote. We determined 500 simulations was sufficient to reach a stability asymptote. **o)** We next assessed the distribution of pseudotime by real age (E11.5 to P4). There is a general progression across pseudotime with age with overlap between stages, as expected. **p)** All 64,004 downsampled cells run in this URD analysis are displayed as a UMAP plot colored by pseudotime value. **q)** UMAP plot from (p) colored and labeled by segment. **r)** URD dendrogram of the 8 general populations colored by segment, as in **Fig. 1f,g. s-z**) Protein expression level changes across each pseudotime trajectory for all proteins in the DRG panel.

**Extended Data Fig. 4.**
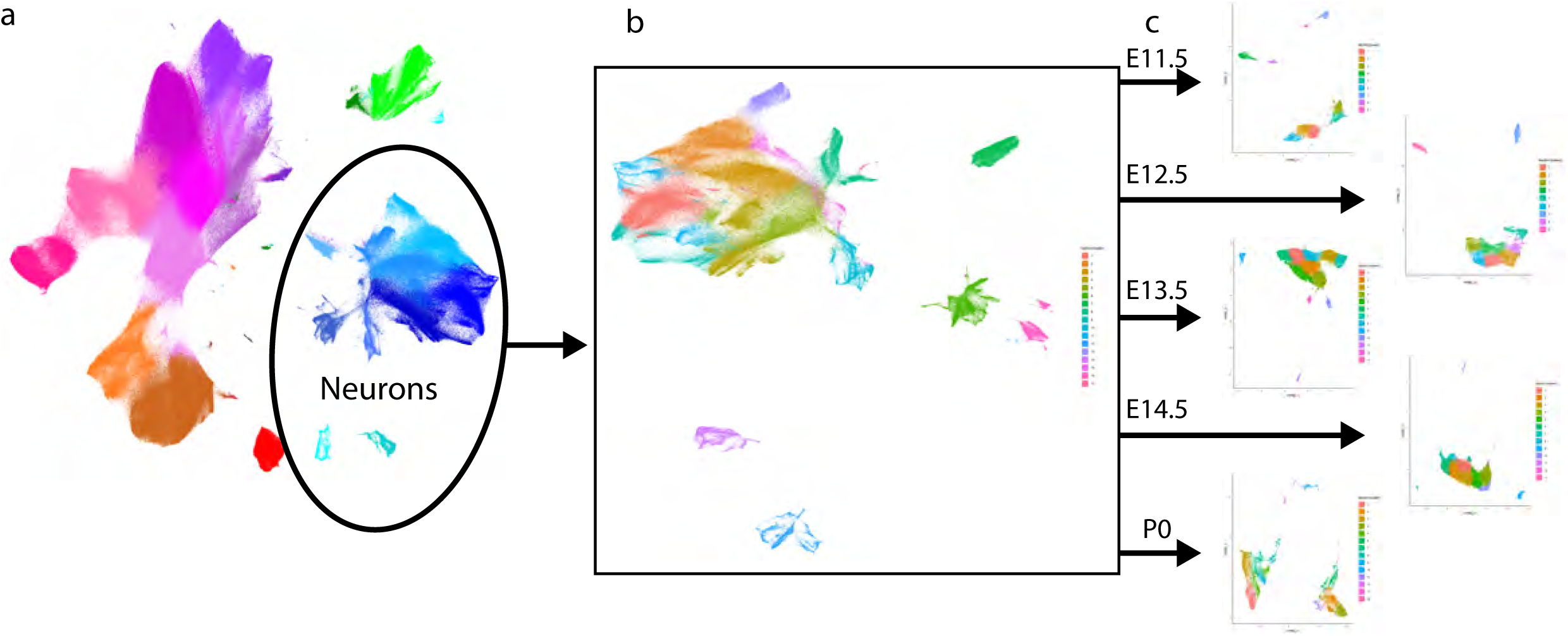
Extraction of single age neuron sets for comparison to IHC. **a)** Neuronal clusters were extracted from the DRG time course (Fig. 1c), **b)** and then subjected to secondary Leiden clustering and UMAP embedding. **c)** Then, individual ages matched to IHC samples were further extracted and subjected to tertiary Leiden clustering and UMAP embedding.

**Extended Data Fig. 5.**
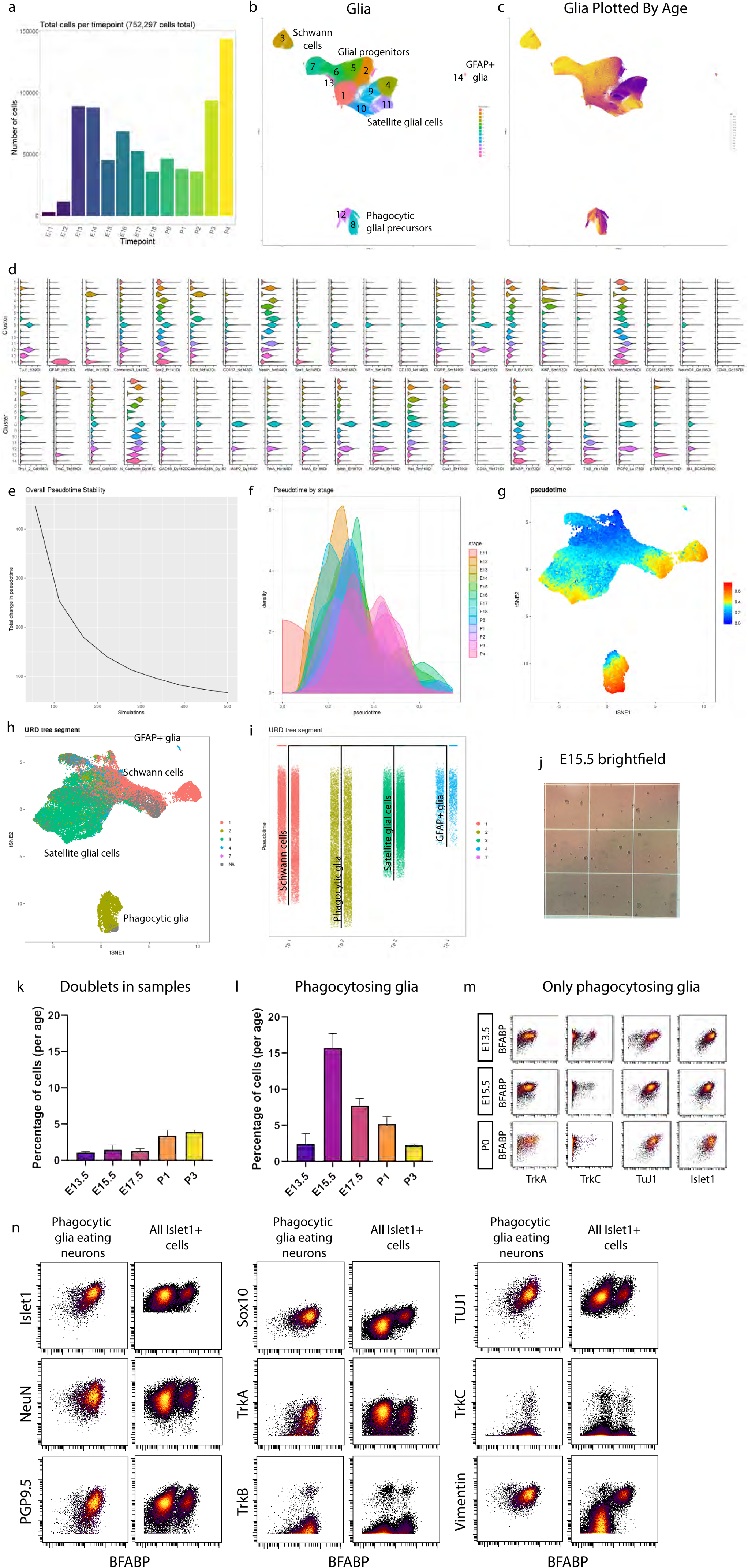
High dimensional analysis of all glial cells and precursors. **a)** Cell number per age for all glia and glial precursors. **b)** Unmodified UMAP embedding from Fig. 4a. White space was trimmed for ease of visualization. **c)** UMAP plot from (b) colored by age. **d)** Violin plot of all markers for the glial clusters (from Fig. 3a) **e-i)** Supplemental URD analysis, performed as in **Extended Data** Fig. 4: **e)** pseudotime stability to calculate simulation number, **f)** pseudotime by stage, **g)** UMAP plot colored by pseudotime value for all 63,796 downsampled cells included in this analysis, **h)** UMAP plot from (g) colored by URD segment, **i)** URD dendrogram colored by segment. **j)** Brightfield image of E15.5 sample on a hemocytometer. **k)** Percentage of doublets counted in the samples from ages E13.5, E15.5, E17.5, P1, and P3 from manual inspection under brightfield analysis with hemocytometer. **l)** Percentage of phagocytic glia by mass cytometry after all cleanup gating. **m)** Biaxial scatterplots from only putative phagocytic glia showing expression of 4 neuronal markers (TrkA,TrkC, TuJ1, and Islet1), by satellite glial cell marker BFABP. **n)** Biaxial scatterplots comparing marker expression for several neuronally expressed markers (Islet1, NeuN, PGP9.5, TuJ1, TrkA, TrkB, and TrkC) and glial markers (Sox10, TrkB, and Vimentin) by BFABP between only putative phagocytic glia and all Islet1^+^ cells (neurons and putative phagocytice glia).

**Extended Data Fig. 6.**
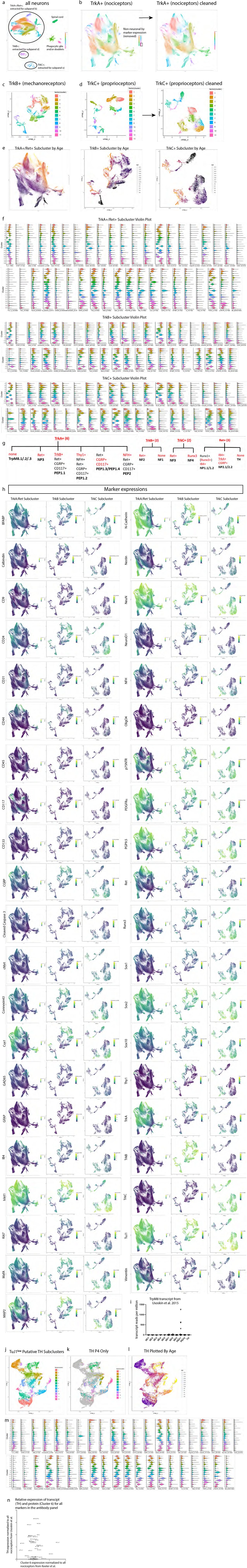
High dimensional analysis of all neurons. **a)** Leiden clustering and UMAP embedding of neurons extracted from the whole time course, labeled by cell type. Circles indicate the 3 main neuronal subtypes by RTK expression: TrkA^+^;Ret^+^, TrkB^+^, and TrkC^+^, respectively. **b)** Leiden clustering and UMAP embedding of the TrkA^+^;Ret^+^ neurons, extracted from a). Three small clusters that did not exhibit neuronal markers were removed from the dataset before a final round of Leiden clustering and UMAP embedding (plot on right). **c)** Leiden clustering and UMAP embedding of the TrkB^+^ neurons extracted from a). **d)** Leiden clustering and UMAP embedding of the TrkC^+^ neurons extracted from a). Putative phagocytic glia expressing TrkC^+^ could not be removed from TrkC^+^ neurons in previous analytic iterations, but this could be done at this resolution resulting in a “cleaned” TrkC^+^ neuronal clustering and UMAP embedding (plot on right). **e)** TrkA^+^;Ret^+^, TrkB^+^, and TrkC^+^ UMAP plots colored by age. **f)** Violin plots of all markers for TrkA^+^;Ret^+^, TrkB^+^, and TrkC^+^ neurons. **g)** Key markers in our panel that allow identification of somatosensory DRG populations^5^. **h)** UMAP plots colored by expression for all panel markers for TrkA^+^;Ret^+^, TrkB^+^, and TrkC^+^ neurons. **i)** TrpM8 transcript data from Usoskin et al., 2015 showing that TrpM8-expressing neurons are a subset of peptidergic nociceptors. **j)** UMAP plot of TH^+^ cells that were extracted from Fig. 4a (Cluster 6) and reclustered. **k)** UMAP plot of the TH^+^ cells from P4 overlaid on the grayed out UMAP plot from j). These were the cells used in the comparison to the Usoskin et al., 2015 transcript data in Fig. 4 **i-n**. **l)** UMAP plot from j) colored by age. **m)** Violin plot of marker expression of all clusters from j). **n)** Comparison of TH^+^ C-LTMR transcripts expression to protein expression for all markers in the mass cytometry panel^2^. In both cases, transcript or protein expression in C-LTMRs was normalized to all nociceptors. However, these normalized expressions are not precisely comparable due to the difference in expression; most mass cytometry expression has a low but non-zero background.

**Extended Data Fig. 7.**
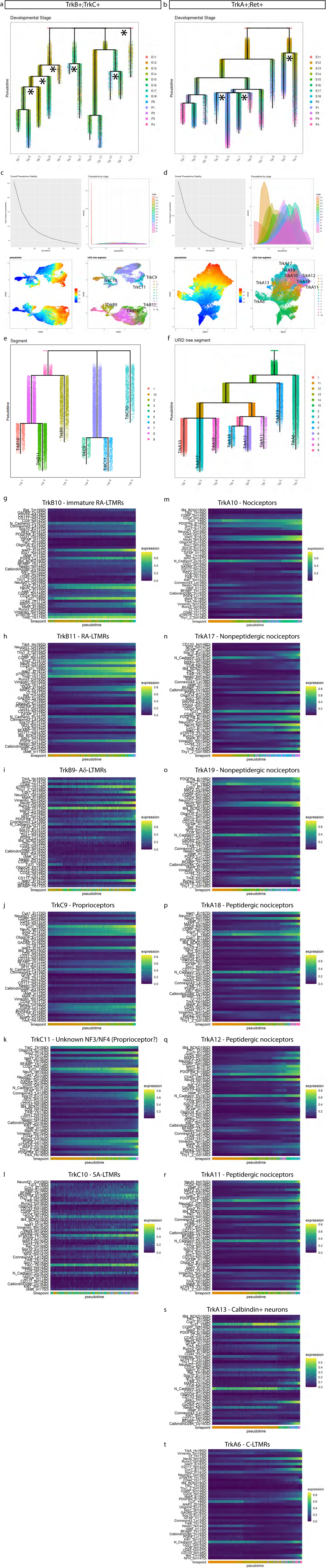
URD pseudotime analysis of all neurons. **a,b)** We initially defined tips as clusters present with ≥ 1% at P4 and all clusters at E11.5 as the root. However, this produced excessively branched pseudotime dendrograms for both a) TrkB^+^;TrkC^+^ and b) TrkA^+^;Ret^+^ where intermediate and more mature cell states were paired as tips. For instance, TrkA^+^/Ret^+^ clusters 5 and 13 are presumptively immature nonpeptidergic nociceptors. The presence of these immature cell types at P4 is expected as cell populations mature over development. By removing presumptive intermediates as tips, we were able to produce the most appropriate molecular trajectory across pseudotime, such as Fig.7 **b-e**. **c-f)** Supplemental URD analysis, same as in **Extended Data** Fig. 3: **c,d)** pseudotime stability to calculate simulation number, pseudotime by stage, UMAP of pseudotime value for all 39,944 (TrkB^+^;TrkC^+^) not downsampled and 64,997 (TrkA^+^/Ret^+^) downsampled cells included in this analysis, UMAP colored by URD segment, for TrkB^+^;TrkC^+^ and TrkA^+^/Ret^+^ datasets, respectively, and **e,f)** URD dendrogram colored by URD segment for TrkB^+^;TrkC^+^ and TrkA^+^/Ret^+^ datasets, respectively. **g-t)** Heatmaps for all URD tips from TrkB^+^;TrkC^+^ and TrkA^+^/Ret^+^ datasets, respectively, for all markers.

**Extended Data Fig. 8.**
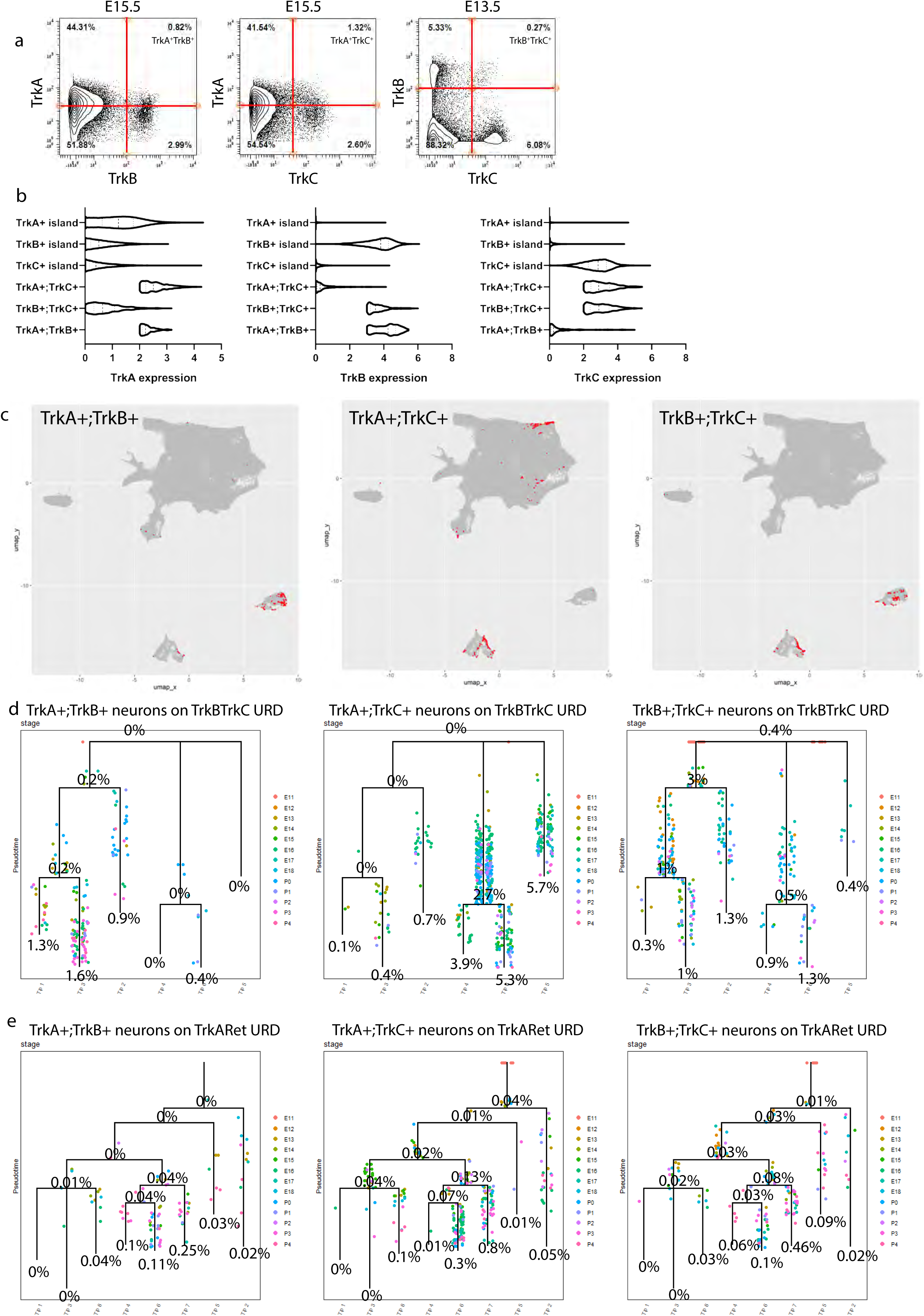
Double Trk^+^ neurons exhibit altered protein expression. **a)** Biaxial scatterplots showing the neurons that express at least two Trks for all 3 combinations: TrkA;TrkB, TrkA;TrkC, and TrkB;TrkC. **b)** Violin plots of Trk expression intensity for all neurons expressing a single Trk (Fig. 4 **a-c**) compared to double-Trk^+^ neurons. **c)** Grayed out UMAPs of all neurons with the neurons expressing multiple Trks highlighted in red. These neurons form subclusters within existing clusters that our analytic pipeline is unable to identify as distinct communities due to their scarcity. **d,e)** URD dendrograms from Fig. 5a without the cells from the URD dataset. Instead, neurons expressing each combination of two Trks (TrkA and TrkB, TrkA and TrkC, and TrkB and TrkC) are mapped on the URDs, colored by the age of each cell, over the TrkB;TrkC URD (d) and the TrkA;Ret URD (e). The numbers of these double-expressors was counted and compared to the number of cells within the URD segment. Since all the TrkB^+^ and TrkC^+^ neurons were added to the URD, this proportion is reported over the URD dendrograms (d). However, the TrkA^+^ and Ret^+^ neurons were downsampled (64,997 out of 492,982 neurons). Thus we multiplied the number of neurons in each segment of the TrkA;Ret URD by the downsampling coefficient (7.584689) and then determined the proportion of the double-expressors over the URD (e).

**Extended Data Fig. 9.**
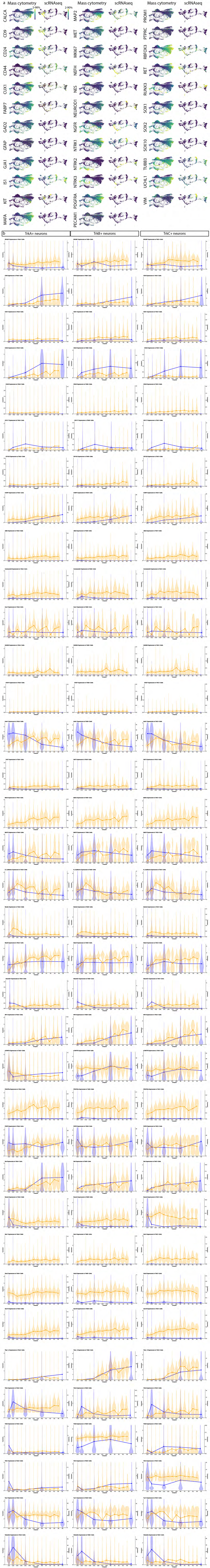
Mass cytometry and scRNA-seq comparison. **a)** FLOW-MAP plots from Fig. 7b-c, colored by each protein marker in the DRG antibody panel for comparison between the mass cytometry and the scRNA-seq^19^. In cases where expression was not detected in the scRNA-seq dataset, the FLOW-MAPs are colored a single intermediate color (e.g. as in GAD2, GFAP, etc). Markers are named with the gene name. **b)** Graphed expression levels for all markers in the protein panel for TrkA^+^, TrkB^+^, and TrkC^+^ populations compared between mass cytometry and scRNA-seq.

**Supplementary Table 1 Antibody panel tailored for dorsal root ganglia development.** Each antibody or reagent used in the mass cytometry staining panel is listed with antigen, conjugated metal, manufacturer, product number, antibody clone, and concentration. Antibodies recognizing surface proteins were stained prior to methanol permeabilization in a single master mix and intracellular proteins, including nuclear proteins, were stained in a single master mix post-permeabilization, denoted by surface or intracellular stain protocol, respectively.

**Supplementary Table 2 Sample data.** Every sample included in this study is reported. For each sample, this includes the age of the mice, the date the cells were collected, the genotype, number of animals in the sample, sex for postnatal samples (all prenatal litters were not sex separated), the total cell count, and which mass cytometry run “set” the sample belonged to. Color indicates barcode set (blue = barcode set 1, green = barcode set 2, red = barcode set 3).

**Supplementary Table 3 Efficiency across data clean-up.** Data event counts for each sample across the 3 mass cytometry barcode sets for the full time course tracked from mass cytometry reads through preprocessing gating clean-up. Data events were first analyzed for the first two barcode sets, each comprising one replicate sample set of a complete time course from E11.5 to P4. These samples are first gated by barcode stringency to ensure that samples without a clear and clean barcode label are removed. Then data events outside of normal parameters for a mass cytometry read are gated out. The remaining events are gated by the viability dye cisplatin and DNA intercalator Iridium to remove damaged cells or non-single cells. Finally, empty isotope channels are gated to remove events with excessive background to produce a population of cleaned data entitled Cells to Analyze. An Islet1^+^ gate enabled an estimate on the percentage of neurons at each age. Using these initial numbers we determined a handful of ages to collect additional samples (barcode set 3). Barcode set 3 was also analyzed across gating clean-up.

